# Cobolt: Joint analysis of multimodal single-cell sequencing data

**DOI:** 10.1101/2021.04.03.438329

**Authors:** Boying Gong, Yun Zhou, Elizabeth Purdom

## Abstract

A growing number of single-cell sequencing platforms enable joint profiling of multiple omics from the same cells. We present Cobolt, a novel method that not only allows for analyzing the data from joint-modality platforms, but provides a coherent framework for the integration of multiple datasets measured on different modalities. We demonstrate its performance on multi-modality data of gene expression and chromatin accessibility and illustrate the integration abilities of Cobolt by jointly analyzing this multi-modality data with single-cell RNA-seq and ATAC-seq datasets.

## 1 Background

Single-cell sequencing allows for quantifying molecular traits at the single-cell level, and there exist a wide variety of platforms that extend traditional bulk platforms, such as mRNA-seq and ATAC-seq, to the single cell. Comparison of different cellular features or *modalities* from cells from the same biological system gives the potential for a holistic understanding of the system. Most single-cell technologies require different cells as input to the platform, and therefore there remains the challenge of linking together the biological signal from the different modalities, with several computational methods proposed to estimate the linkage between the different modalities, such as LIGER [Welch et al., 2019] and Signac (Seurat) [Stuart et al., 2020, 2019].

Recently, there are a growing number of platforms that allow for measuring several modalities on a single cell. CITE-seq [Stoeckius et al., 2017] jointly sequences epitope and transcriptome; scNMT-seq [Clark et al., 2018] jointly profiles chromatin accessibility, DNA methylation, and gene expression; and sci-CAR [Cao et al., 2018], Paired-seq [Zhu et al., 2019], and SNARE-seq [Chen et al., 2019] enable simultaneous measurement of transcription and chromatin accessibility (we direct readers to Lee et al. [2020] for a comprehensive review). By directly measuring the different modalities on the same cells, these techniques greatly enhance the ability to relate the different modalities. With the emergence of joint platforms, new computational methodologies for analyzing multi-modality data have also been developed. Early methods mainly focused on CITE-seq [Gayoso et al., 2021, Wang et al., 2020, Hao et al., 2020], which jointly sequences gene expressions and at most a few hundred antibodies. Recently, more methods have been proposed to enable the modeling of cells with simultaneous measurement of gene expression and higher-dimensional modalities such as chromatin accessibility, such as MOFA+ (also known as MOFA2 Argelaguet et al. [2020]), scMVAE Zuo and Chen [2021], BABEL [Wu et al., 2021], and scMM [Minoura et al., 2021].

However, single-cell datasets on a single modality are far more common, and are usually of higher throughput. Indeed it is natural that joint-modality data from the same system will be used to augment single-modality data, or vice versa. Therefore, there is a critical need for an analysis tool that is both a stand-alone application for multi-modality data as well as a tool for integration of these datasets with single-modality platforms. BABEL [Wu et al., 2021] and scMM [Minoura et al., 2021], while not directly targeting this task, do allow the use of the joint-modality data to predict one single-modality dataset into another type of modality. However, neither directly integrate the data together to allow for downstream analysis of the joint set of data regardless of modality, such as cell subtype detection.

Our method `Cobolt` fills this gap by providing a coherent framework for a full integrative analysis of multi-modality and single-modality platforms. The result of `Cobolt` is a single representation of the cells irrespective of modalities, which can then be used directly by downstream analyses, such as joint clustering of cells across modalities. `Cobolt` estimates this joint representation via a novel application of Multimodal Variational Autoencoder (MVAE) [Wu and Goodman, 2018] to a hierarchical generative model. The integration of the single-modalities is done by a transfer learning approach which harnesses the valuable information found by joint sequencing of the same cells and extends it to the cells in the single-cell platform. The end result is a single representation of all of the input cells, whether sequenced on a multi-modality platform or a single-modality platform. In this context, `Cobolt` gives an over-all integrative framework that is flexible for a wide range of modalities.

We demonstrate `Cobolt` on two use-cases. The first uses `Cobolt` to analyze only a multi-modality sequencing dataset from the SNARE-seq technology; we show that `Cobolt` provides a joint analysis that better distinguishes important facets of each modality, compared to existing methods. The second demonstrates the use of `Cobolt` to integrate multi-modality data with single-modality data collected from related biological systems, where `Cobolt` creates a joint representation that can be used for downstream analysis to provide meaningful biological insights. We show that `Cobolt` also performs better than related tools in this integrative task.

## 2 Results

### 2.1 The Cobolt Model

We develop a novel method, `Cobolt`, that utilizes joint-modality data to enable the joint analysis of cells sequenced on separate sequencing modalities. We do this by developing a Multimodal Variational Autoen-coder based on a hierarchical bayesian generative model. We briefly describe the premise of the model using the example of two modalities: mRNA-seq and ATAC-seq (for more details in greater generality, see the Methods Section 5). We assume that we have a set of cells with both mRNA-seq and ATAC-seq data collected using the joint modality platform (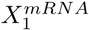 and 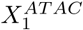), as well as (optionally) a set of cells with only mRNA-seq data 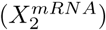, and a set of cells with only ATAC-seq data 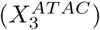. `Cobolt` takes all of this data as input to find a representation of the cells in a shared reduced dimensionality space, regardless of modality (Figure 1A).

**Figure 1:**
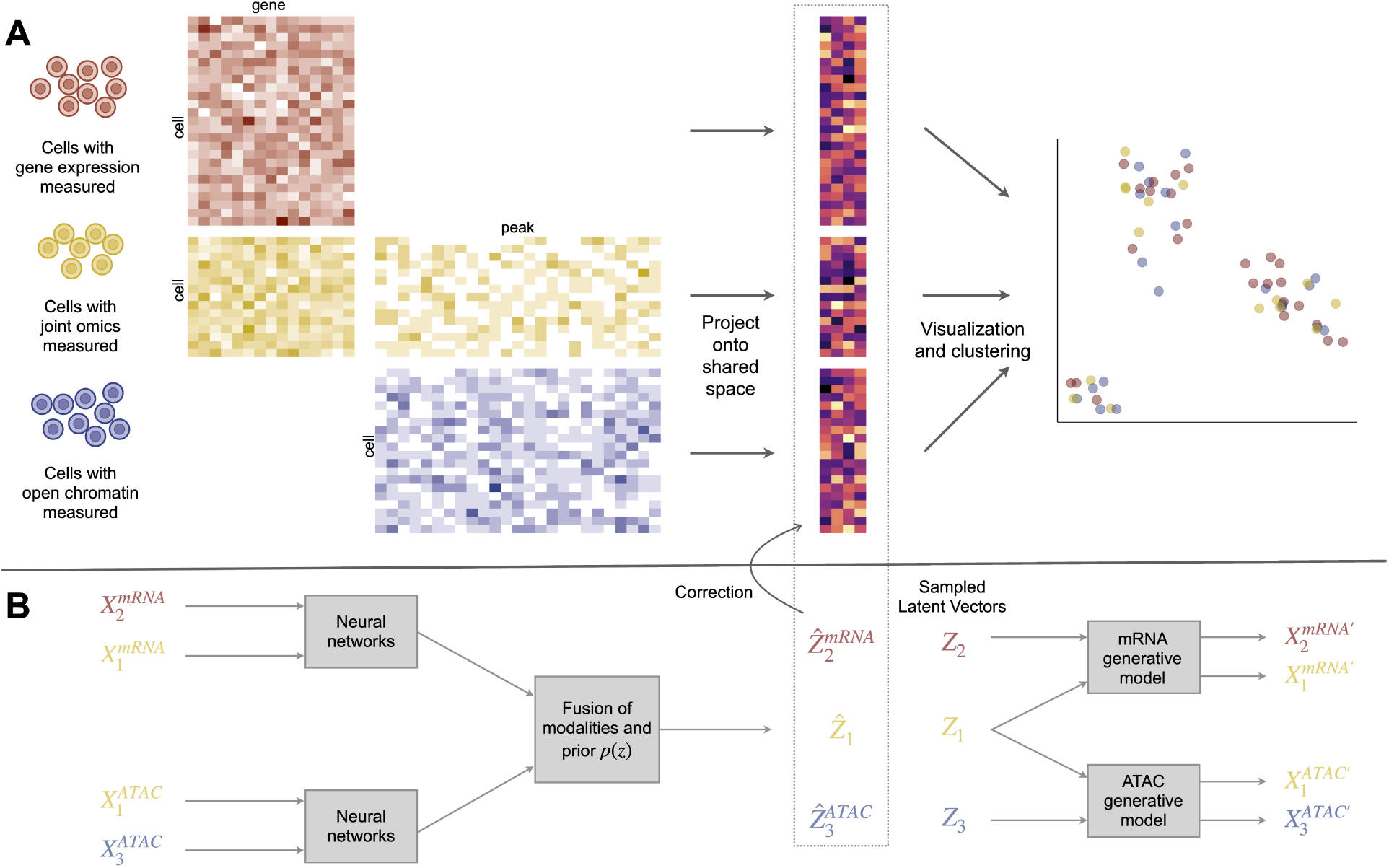
An overview of the Cobolt method. The upper panel shows the workflow of `Cobolt`, which takes as input datasets with varying modalities, projects the data into shared latent space, and then performs visualization and clustering with all datasets combined. The lower panel shows the `Cobolt` variational autoencoder model with encoders plotted on the left and decoders on the right.

`Cobolt` models the sequence counts from the modalities inspired by Latent Dirichlet Allocation (LDA) [Blei et al., 2003] model. The LDA model is a popular Bayesian model for count data which has been successfully applied to genomics in areas such as clustering of single-cell RNA-seq data [Yotsukura et al., 2016, Sun et al., 2018], single-cell ATAC-seq analysis [González-Blas et al., 2019], and functional annotation [Backenroth et al., 2018]. `Cobolt` builds a hierarchical latent model to model data from different modalities and then adapts the MVAE approach to both estimate the model and allow for a transfer of learning between the joint-modality data and the single-modality data.

`Cobolt` assumes that there are *K* different types of possible categories that make up the workings of a cell. For ease of understanding, it is useful to think of these categories as biological processes of the cell, though the categories are unlikely to actually have a one-to-one mapping with biological processes. Each category will result in different distributions of features in each of the modalities – i.e., different regions of open chromatin in ATAC-seq or expression levels of genes in mRNA-seq for different categories. The features measured in a cell are then the cumulative contribution of the degree of activation of each category present in that cell. The activation level of each category is represented by the latent variable *θ*_*c*_ for each cell *c*, which gives the relative activity of each of the *K* categories in the cell. *θ*_*c*_ is assumed to be an intrinsic property of each cell representing the underlying biological properties of the cell, while the differences of data observed in each modality for the same cell are due to the fact that the categories active in a cell have different impacts in the modality measured (open chromatin in ATAC-seq versus gene expression in mRNA-seq). We assume *θ*_*c*_ = *σ*(*z*_*c*_), where *z*_*c*_ is drawn from a Gaussian prior and *σ* is the soft-max transformation; this is an approximation to the standard Dirichlet prior for *θ*_*c*_ that allows use of variational autoencoders to fit the model [Srivastava and Sutton, 2017]. The mean of the posterior distribution gives us an estimate of our latent variable *z*_*c*_ for each cell, and the posterior distribution is estimated using variational autoencoders (VAE).

In the end, `Cobolt` results in an estimate of the latent variable *z*_*c*_ for each cell, which is a vector that lies in a *K*-dimensional space. This space represents the shared biological signal of the individual cells, irregardless of modality, and can be used for the common analysis tasks of single-cell data, such as visualization and clustering of cells to find subtypes. Importantly, we can predict the latent variable *z*_*c*_ even when a cell does not have all modalities measured. Moreover, because of the joint-modality platforms, `Cobolt` does not require that the different modalities measure the same features in order to link the modalities together – the fact that some of the cells were sequenced on both platforms provides the link between different types of features. Therefore, ATAC-seq peaks and mRNA-seq gene expression can be directly provided as input. This is unlike methods that do not make use of the joint-modality data and require that the different modalities be summarized on the same set of features, for example by simplifying ATAC-seq peaks to a single measurement per gene.

### 2.2 Cobolt as an analysis tool for multi-modality data

While the full power of `Cobolt` is to integrate together data from single and multiple modality datasets, in its simplest form `Cobolt` can be used for the analysis of data from solely a multi-modality technology. We demonstrate this usage with the SNARE-seq data [Chen et al., 2019], which consists of paired transcription and chromatin accessibility sequenced on 10,309 cells of adult mouse cerebral cortices.

We first compare Cobolt with a simple, but common, approach for analyzing joint-modality data: the two modalities are analyzed separately and then the results are linked together. This is the strategy of [Chen et al., 2019], where the authors primarily clustered the gene expression modality to form clusters of the cells, and then performed a separate analysis on chromatin accessibility modality as a comparison. Focusing on the gene expression modality is common, since it is often assumed to have the greatest resolution in determining cell-types. However, the reduced representations and clusters created on one modality may not be representative of all the underlying cell subtypes. Indeed, when we perform clustering analysis on only the gene expression modality using Seurat [Stuart et al., 2019] and only the chromatin accessibility modality using cisTopic [González-Blas et al., 2019] (consistent with [Chen et al., 2019], see Section 5.3 for details), both modalities find distinct clusters that are not reflected in the other modality (Supplementary Figure 1, Additional File 1). For example, cells identified by marker genes as non-neuronal cells, such as astrocytes, oligodendrocytes, oligodendrocyte precursors, and microglial cells are clustered into their respective cell-types based on mRNA-expression but are not separated based only on chromatin accessibility (Supplementary Figure 2A, Additional File 1). Similarly, a subset of Layer 5/6 cells has distinct chromatin accessibility peaks but are intermingled with other Layer 5/6 cells in the gene expression clusters; these differential peaks include one near the gene *Car12* which is a marker gene of the previously annotated subtype of Layer 6 (L6 Car12, Tasic et al. [2016]) and which shows higher expression in this subset of cells (Supplementary Figure 2BC, Additional File 1). A joint analysis with `Cobolt`, unlike the single-modality analyses, finds these subtypes detected by only one modality and not the other (Supplementary Figure 1, Additional File 1).

Next we compare with other methods that explicitly analyze the two modalities jointly, like `Cobolt`. We consider the methods MOFA2, scMM, and BABEL. MOFA2 uses Bayesian group factor analysis for dimensionality reduction of multi-modality datasets; BABEL trains an interoperable neural network model on the paired data that translates data from one modality to the other; and scMM [Minoura et al., 2021] uses a deep generative model for joint representation learning and cross-modal generation. We apply each of these methods to the SNARE-seq data.

Since a joint analysis method should be able to reflect subtype signals captured by all modalities, we similarly evaluate the methods on how well their lower-dimensional spaces represent separate clusters identified separately in each modality, as described above. In Figure 2, we visualize the lower-dimensional space generated by `Cobolt`, MOFA2, scMM, and BABEL via UMAP (Uniform Manifold Approximation and Projection [McInnes et al., 2018]), where we color the cells based on the clusters found by clustering the gene expression modality data (Figure 2A) and those based on clustering the chromatin accessibility data (Figure 2B). We would note that BABEL does not create a single reduced-dimensionality representation for a paired cell, but rather one per modality (the two latent representations are learned jointly and are quite similar). BABEL’s lower-dimensionality representation does quite poorly, separating major clusters of cells found in both modalities, such as layer 2 to 6 intratelencephalic (IT) neurons (colored red, purple, pink, and cyan in Figure 2A). Both MOFA2 and scMM capture these large clusters, which are shared between the modalities. However, we see clusters specific to a single modality not reflected on their lower-dimensional space. For example, the highlighted gene expression cluster in Figure 2A is practicably indistinguishable in the scMM UMAP but separated in the `Cobolt` UMAP. Differential mRNA expression analysis between this cluster and neighboring cells finds strong expression of known markers of Layer 6 cells in this cluster (*Col24a1* Baker et al. [2018], *Gnb4* Sorensen et al. [2015], *Rxfp1* Belgard et al. [2011], *Nr4a2* Zeng et al. [2012], Fazel Darbandi et al. [2018], and *Ntng2* Zeng et al. [2012], Supplementary Figure 4A, Additional File 1) as well as strong expression of *Car3* defined in Yao et al. [2021] as a marker of a subset of Layer 6 IT cells. Neighboring cells do not express these known marker genes and instead express layer 5/6 IT markers (cyan cluster) or layer 2/3 IT markers (red cluster), indicating that this cluster missed by scMM consists of a biologically meaningful subset of Layer 6 IT cells. Similarly, in Figure 2B, we highlight two clusters of cells which clearly separate in the `Cobolt` analysis and are separate clusters based on both mRNA-Seq and chromatin profiles, but are mixed together in the MOFA2 analysis. Differential mRNA expression analysis between these clusters reveal genes *Adarb2* and *Sox6* differ in expression between these groups (Supplementary Figure 4B, Additional File 1), which are known markers whose expression distinguish the CGE and Pvalb clusters respectively Yao et al. [2021]. Integrative analysis in Section 2.3 confirms this identification by integrating this SNARE-Seq data with annotated scRNA-Seq data and placing these cells with cells annotated as CGE and Pvalb in Yao et al. [2021].

**Figure 2:**
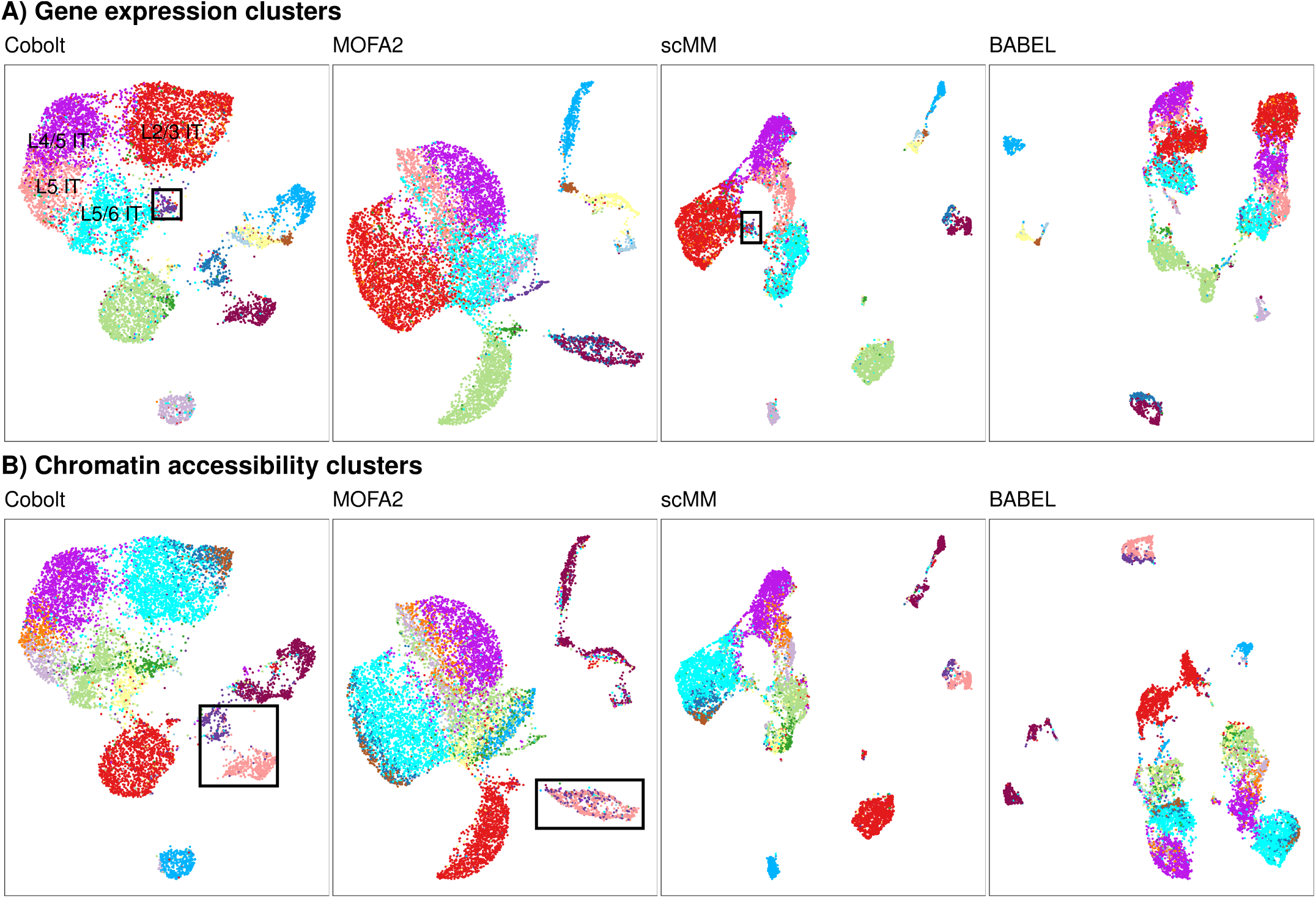
Comparison of multi-modality analysis methods. A, B) UMAP visualizations of the reduced dimensionality space created by `Cobolt`, MOFA2, scMM, and BABEL. The cells are color-coded by the cluster they are assigned to based on clustering of A) only gene expression modality and B) only chromatin accessibility modality. We note that the cluster colors are randomly and separately assigned for panel A and B. Highlighted in the panels are clusters that are well separated in the analysis of `Cobolt`, but not the other methods. C) Mean silhouette width of the gene expression clusters and chromatin accessibility clusters evaluated on the UMAP representations generated by `Cobolt`, MOFA2, scMM, and BABEL. More details on the silhouettes per cluster can be found in Supplementary Figure 3 (Additional File 1).

To quantitatively evaluate these observations, we calculate the average silhouette widths of the modality-specific clustering on the UMAPs generated by `Cobolt`, MOFA2, scMM, and BABEL (shown in Figure 2C), where higher silhouette widths indicate that cells are closer to other cells in the same cluster. As expected from our observations, BABEL’s representation results in extremely small silhouette widths, reflecting the many clusters separated in BABEL’s representation. MOFA2 has the smallest silhouette width on chromatin accessibility clusters, supporting our observation that its joint space does not represent this modality well; similarly, scMM gives relatively small measures on the gene expression modality. `Cobolt` best represents both modalities with the highest silhouette width measure.

### 2.3 Cobolt for integrating multi-modality data with single-modality data

We now turn to integrating multi-modality data with single-modality data. For this use-case, we use `Cobolt` to jointly model three different datasets — the SNARE-seq of mouse cerebral cortices analyzed in the above section, together with a scRNA-seq and a scATAC-seq dataset of mouse primary motor cortex (MOp) [Yao et al., 2021]. In addition, we also demonstrate `Cobolt` for joint modelling of single-cell sequencing of human peripheral blood mononuclear cells (PBMCs): two multi-modality datasets pairing ATAC and mRNA measurements on 10,970 and 12,012 cells from different samples of the 10X Multiome platform 10x Genomics [2021, August 9c,,], combined with 23,837 cells from scRNA-Seq 10x Genomics [2021, August 9b] and 9,030 cells from scATAC-Seq 10x Genomics [2021, August 9a].

The result in both examples is a lower-dimensional latent space that aligns the different modality data into a single representation. In Figure 3A and B, we visualize this low-dimensional space via UMAP for the mouse cortex and human PBMCs, respectively, with cells colored by their data set of origin. We see that the cells from different datasets are well aligned regardless of their source of origin.

**Figure 3:**
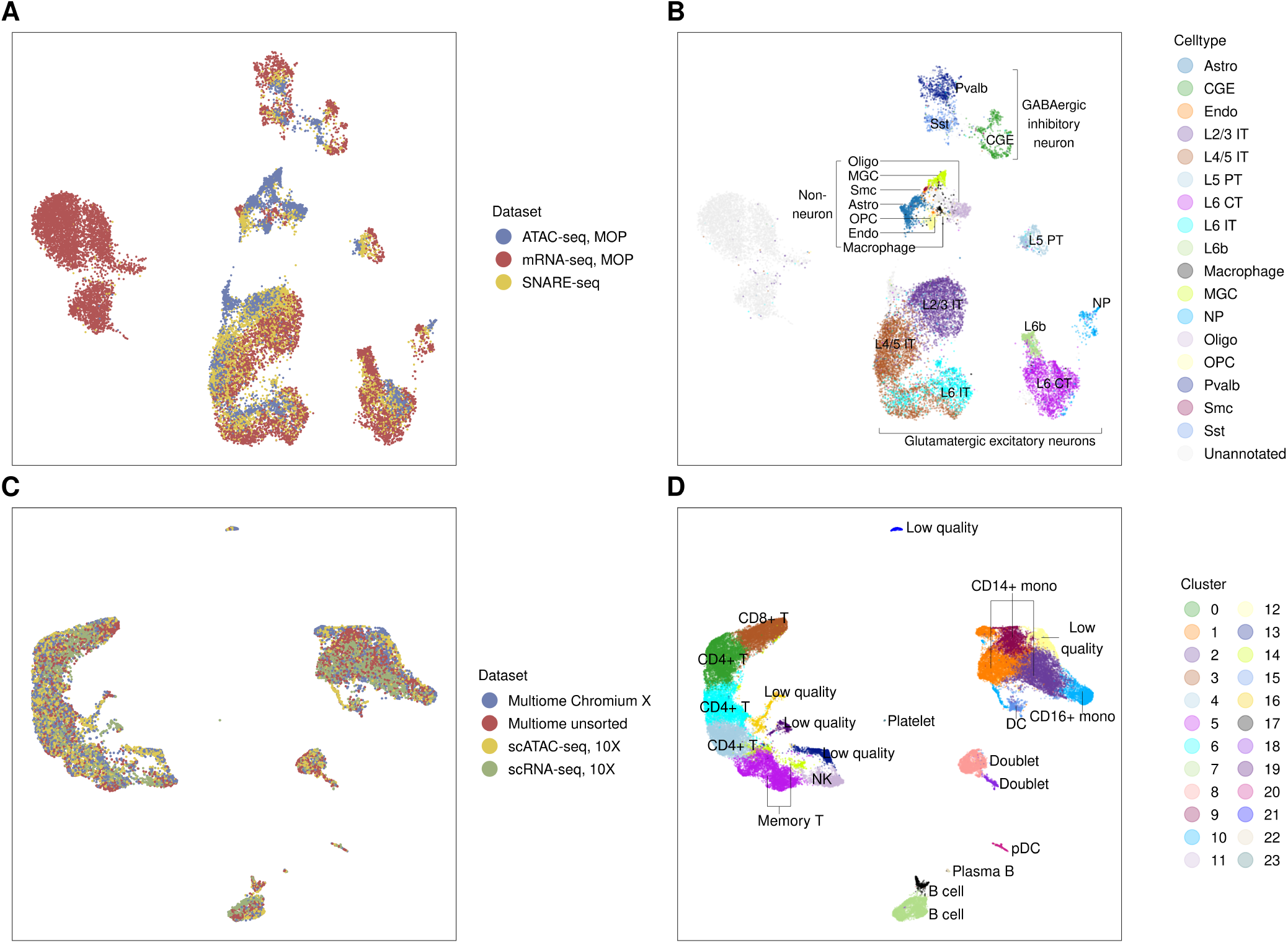
The UMAP visualization of A, B) the mouse cortex integration and C, D) the 10X PBMC integration. Cells are colored in A, C) by dataset of origin, in B) known cell type annotation of [Yao et al., 2021], and in D) by our *de novo* clustering and annotated based on gene markers. For the mouse cortex integration, both the MOp scRNA-seq and the MOp scATAC-seq contain a substantial fraction of cells labeled “unannotated” by the authors of the data and that do not map to known cell types. The cell type abbreviation largely follows the data paper [Yao et al., 2021]: astrocytes (Astro); caudal ganglionic eminence interneurons (CGE); endothelial cells (Endo); layer 2 to layer 6 (L2-6); intratelencephalic neurons (IT); pyramidal tracts (PT); corticothalamic neurons (CT); L6b excitatory neurons (L6b); microglial cells (MGC); near-projecting excitatory neurons (NP); oligodendrocytes (Oligo); oligodendrocyte precursors (OPC); smooth muscle cells (SMC); medial ganglionic eminence interneurons subclasses based on marker genes (Sst, Pvalb). For the 10X PBMC integration the following abbreviations are observed: dendritic cell (DC); plasmacytoid dendritic cells (pDC); monocytes (mono); natural killer cells (NK).

To consider further the biological meaning of the lower-dimensional representation, we label the MOp cells from the mouse cortex dataset in Figure 3C by their cellular subtype as annotated in [Yao et al., 2021]. For the purposes of comparison across the modalities, we integrated some cell-types in [Yao et al., 2021] into larger groupings and modified the names so as to have comparable groups (see Methods, Section 5.3). For the SNARE-seq cells, we do not have the cell types given in [Chen et al., 2019], so we use the identifications found by our analysis of only the SNARE-seq cells (see Methods, Section 5.3). We see that cells from the same cellular subtypes are projected closely regardless of the data source. We also see that the representation of `Cobolt` respects the larger category of cell types by grouping three major cell classes: GABAergic inhibitory neurons (CGE, Sst, Pvalb), glutamatergic excitatory neurons (IT, L5 PT, L6 CT, L6b, NP), and non-neurons.

The PBMC datasets does not have accompanying annotation, so we applied the Louvain clustering algorithm to the lower-dimensional representation from `Cobolt` to identify potential celltypes. Using marker genes, Ding et al. [2020], Stuart et al. [2019], Pliner et al. [2019], Franzén et al. [2019] we classified the clusters into known subtypes expected for PBMC data (Supplementary Figure 5 and 6, Additional File 1) and we see that important cell types and functions are localized in the `Cobolt` representation (Figure 3C)

The mouse cortex data also demonstrates the ability of our joint representation to capture subtypes signal that are not shared across all of the modalities. Indeed despite detecting mostly similar cell types, the MOp datasets profile several cell types distinct to the modality. For example, microglial cells (MGC) and smooth muscle cells (SMC) are uniquely detected in scATAC-seq. The different datasets also have different cellular compositions of their shared subtypes, where astrocytes (Astro) and oligodendrocytes (Oligo) are much more abundant in the scATAC-seq (6.55% and 10.50%) than in the SNARE-seq (4.5% and 2.84%) and scRNA-seq (0.40% and 0.36%). As shown in Figure 3B, cell populations unique to one dataset are grouped in the UMAP plot and are distinguishable from the other datasets/cell types. This indicates that `Cobolt` reconciles data even when one cell population is entirely absent or scarcely represented in one or more data sources, which is important in integrating datasets collected from related but slightly differing settings.

`Cobolt` also facilitates subtype identification at a finer resolution by transferring information between modalities. For example, for the mouse cortex data, Figure 3B shows the cell types based on the published annotation. To make the annotation consistent between scRNA-Seq and scATAC-Seq, some of these cell types are the result of merging some cell types into larger categories (see Methods, Section 5.3). One cell type, caudal ganglionic eminence interneurons (CGE, dark green in Figure 3B) was annotated in the scATAC-seq MOp dataset as one cluster, but the scRNA-seq annotation further divided CGE cells into 3 subtypes based on marker genes – Lamp5, Vip, and Sncg. Our joint mapping of the cells allows us to relate the subtypes detected in scRNA-seq to scATAC-seq and provides a finer resolution breakdown of CGE in scATAC-seq. Specifically, we ran a *de novo* clustering of our joint mapping of all three datasets by `Cobolt` (see Methods Section 5.3 and Supplementary Figure 7). This clustering results in a cluster (cluster 13) composed of Lamp 5 and Sncg cells, while another (cluster 16) is mostly Vip cells (Figure 4A). This subdivision is further validated by gene expression and gene activity levels in these clusters of marker genes *Lamp5* and *Vip* as well as other genes known to discriminate subtype Lamp5/Sncg from subtype Vip Tasic et al. [2018], such as *Reln* and *Npy* (Figure 4B). We further validated the scATAC-seq clusters through *de novo* differential accessibility (DA) and differential expression (DE) analysis (Supplementary Figure 8). We identified 94 DA genes between these two clusters. 78 of the DA genes are also found DE in the mRNA, and all of them have the same direction of fold changes in the DE and the DA analyses, i.e., the genes with lower/higher gene expression in cluster 13 compared to cluster 16 are also less/more accessible. This shows that the joint model of `Cobolt` can help distinguish noisy cells in one dataset with additional information from other datasets or modalities.

**Figure 4:**
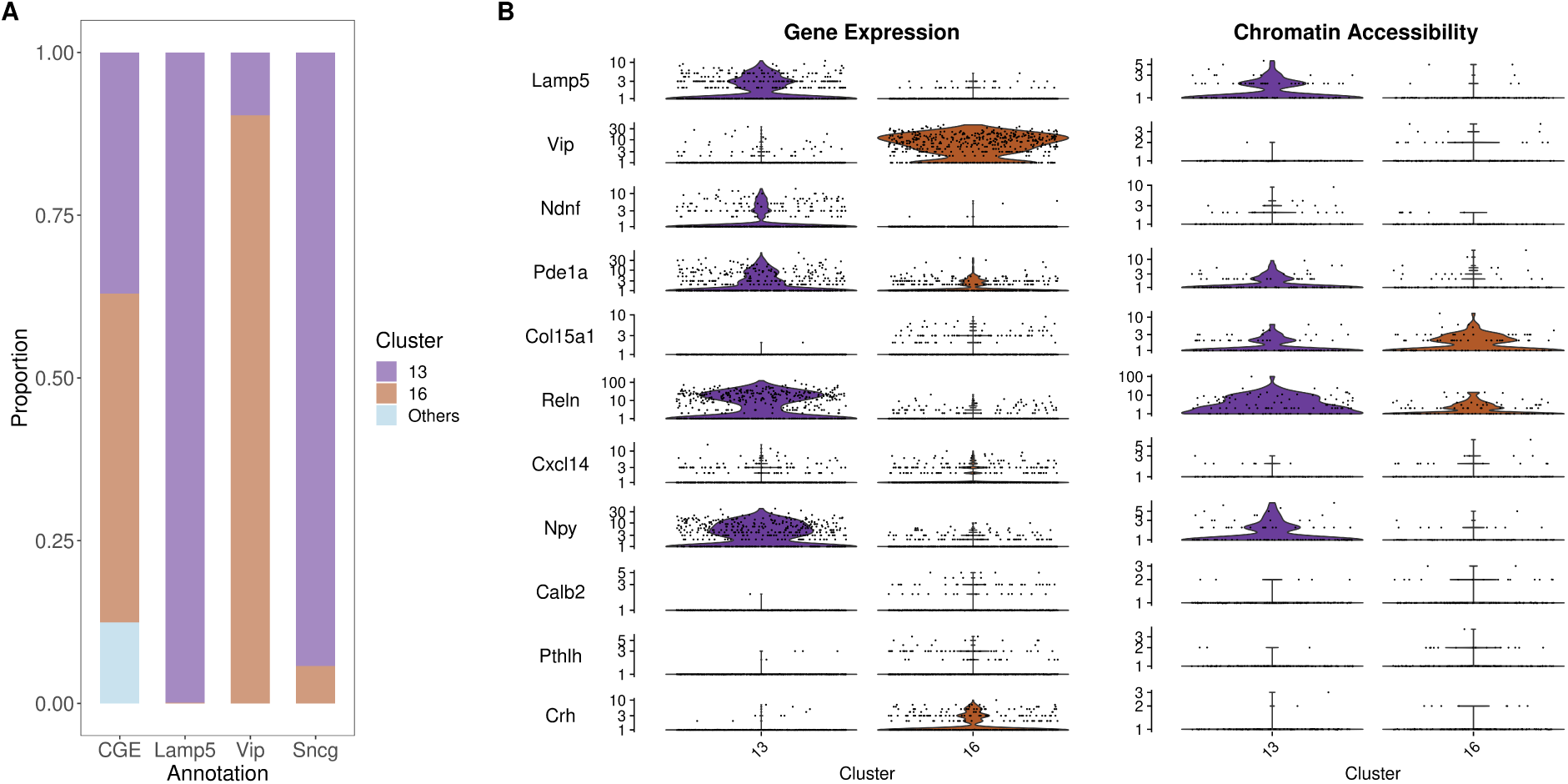
A) We show the relative composition of cells annotated as CGE by the scATAC-seq dataset in the clusters found by clustering of the reduced dimensionality of `Cobolt`, and compare that to the relative composition of the cells annotated in the subtypes Lamp5, Vip, and Sncg by the scRNA-seq dataset. (B) Plots of the gene expression (scRNA-seq) and gene body accessibility summaries (scATAC-seq) in clusters 13 and 16 of the marker genes that distinguish between cell types Lamp5 and Vip [Yao et al., 2021, Tasic et al., 2018].

Furthermore, `Cobolt` is robust to poor-quality cells and low-expressed genes by using a count-based model, which should naturally down-weight the influence of low-count cells. In the above analysis of the mouse cortex data, we filtered out 2.5% of MOp mRNA cells and 15.8% ATAC-seq cells due to low counts and other quality control measures following the work of [Yao et al., 2021]. Yet `Cobolt` is robust to these choices, even in the extreme case where there is no filtering on cells and only a minimal filter on genes is performed (Supplementary Figure 9, Additional File 1).

#### 2.3.1 Comparison with existing methods

As described in the introduction, there are few existing methods that allow full integration of multi-modality sequencing data with single-modality data. MOFA2 only analyzes joint-modality data; BABEL and scMM train a joint model on the paired joint-modality data, and allow the user to apply this model to single-modality datasets to predict the other “missing” modality. Unlike `Cobolt`, these two methods do not use the single-modality data in the training of the model, nor do they provide a representation of the single-modality data in a single shared representation space regardless of modality – for example, for the purposes of joint clustering of all of the cells across modalities.

Therefore, to have additional points of comparison, we apply the LIGER and Signac methods, which are designed for integrating *unpaired* modalities (the implementation details can be found in Section 5.3). LIGER applies an integrative nonnegative matrix factorization (iNMF) approach to project the data onto a lower-dimensional space and then builds a shared nearest neighbor graph for joint clustering. Signac implements canonical correlation analysis (CCA) for dimensionality reduction; Signac subsequently transfers cell labels by identifying mutual nearest neighbor cell pairs across modalities.

To evaluate the performance of these methods when integrating single-modality data with multiple-modality data, we return to the multi-modality datasets described above to create artificial sets of multiple-modality data and single-modality data. For the 10X Multiome data, we make use of the fact that we have two datasets from the 10X Multiome platform run on different patient samples: PBMC of a healthy male donor aged 30-35 (“Multiome Chromium X”) 10x Genomics [2021, August 9c] and PBMC of a female donor aged 25 (“Multiome unsorted”) 10x Genomics [2021, May 3]. We ignore the pairing information in the Multiome unsorted data, and treat the mRNA and ATAC measurements as coming from unpaired, separate sequencing experiments. For the SNARE-Seq data, we randomly assign the cells to be considered as from either the multi-modality dataset (20%) or the single-modality datasets (80%) and run each of the methods. The choice of 20% and 80% was based on the relative size of the SNARE-Seq joint-modality data to the individual MOp scATAC-Seq data and scRNA-Seq datasets, and reflects the fact that single-modality datasets are much higher-throughput than paired-modality data.

We give to each of the methods the multi-modality data from the cells that remain paired, and for the cells where we ignore the pairing information give the mRNA and ATAC data from those cells as if they were single-modality datasets. For LIGER and Signac, which are not designed for multiple-modality data, we hide the pairing information on all cells, and treat all of the cells as if they were collected on different cells. In this way, we have a ground truth on how the cells in the single-modality data sets should be connected to each other, and we can compare the methods by evaluating whether coordinates of the reduced dimensions *Z* for the pairs of cells assigned to the single-modality datasets were close together. Specifically, for each method, we evaluate for each cell in the mRNA single-modality set its coordinates 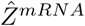 and for a fixed number *k* locate its *k* nearest neighbors in the ATAC single-modality set based on the coordinates 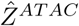. We then calculate the percentage of mRNA single-modality cells whose paired cell in the ATAC single-modality is included in its set of nearest neighbors. The reverse analysis was done using chromatin accessibility as the query and evaluating the percentage whose nearest neighbors include their mRNA pair. Many popular clustering routines use nearest-neighbor graphs for identifying clusters, so this is a metric directly related to whether the cells assigned to the single-modality data would likely correctly cluster together across modalities, but avoids having to specify cluster parameters, especially as applied to different methods (and LIGER and Signac have their own clustering techniques specific for their methods).

As shown in Figure 5, the `Cobolt` joint representation does a much better job for both the SNARE-Seq and 10X multiome data of assigning coordinates to the single-modality cells that place them close to their matching pair. The proportion of single-modality cells that are neighbors to their pair is much larger than any of these other methods in both datasets. Surprisingly, for the SNARE-seq data, the other methods that make use of the joint-modality data to develop their model (scMM and BABEL) do much worse than Signac and LIGER which do not have any information linking the cells together. The 10X multiome data, which has similar numbers of cells from the single-modality datasets as the multi-modality dataset, shows scMM and BABEL perform comparably to Signac, though not as well as Cobolt; similarly we see improved performance of scMM and BABEL in the SNARE-Seq data when we increase the proportion of dual-modality cells to 80% and only 20% of cells being single-modality (Supplementary Figure 12, Additional File 1), but still well below the performance of `Cobolt`. This points to the power of truly integrating the high-throughput single-modality data into the analysis, particularly when there are more cells sequenced from the single-modality data, as is frequently the case.

**Figure 5:**
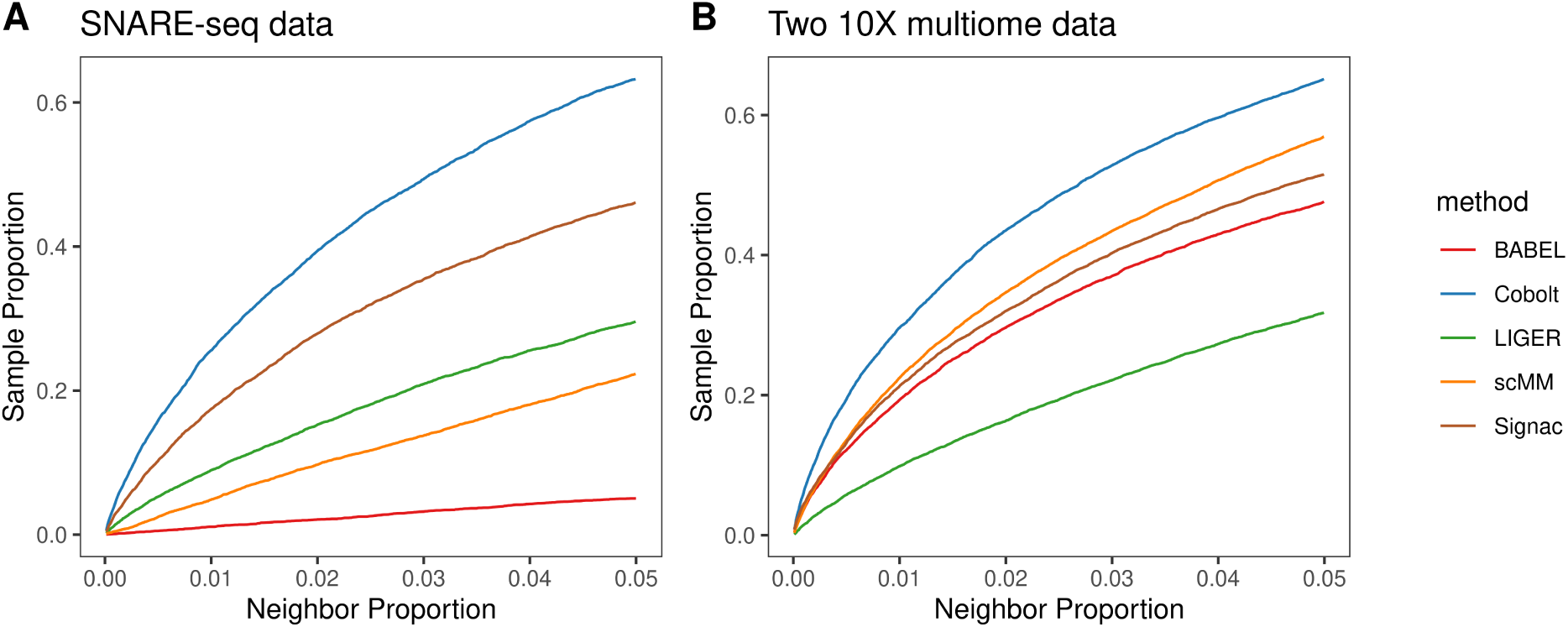
Evaluation of dimensionality reduction. A comparison of lower-dimensional representation generated by LIGER, Signac, scMM, BABEL, and `Cobolt` on A) a SNARE-seq data and B) two 10X multiome datasets. The x-axis shows the number of neighbors considered (*k*) as a proportion of the total testing sample size. The y-axis shows the proportion of cells whose paired data are within their *k*-nearest neighbors in the other modality. The plot gives the results when using gene expression data as the query. We observe very similar results when chromatin accessibility is used as the query (Supplementary Figure 10, Additional File 1).

Subtype detection is a critical aspect of single-cell data analysis, and integrating single-modality data with multiple-modality data gives the potential for higher resolution detection. Our nearest-neighbor metric is directly related to subtype detection, with a higher neighbor proportion corresponding roughly to finding larger clusters in a clustering algorithm. scMM and BABEL only provide cross-modality predictions, rather than a joint embedding of the single-modality data with the multiple-modality data. Downstream tasks such as clustering for subtype detection must be done on either the mRNA expression space or the chromatin expression space, and as we have seen in Section 2.2 each of which can miss important features of the data. Indeed this may be an important factor in their low performance on our nearest-neighbor analysis. LIGER and Signac do provide a joint embedding of all of the data, but do so without the use of the pairing information for the joint-modality data for training the embedding.

The previous analysis provides a comparison with a known ground-truth and the evaluation at different levels of analysis. Now we return to the joint embedding of MOp scRNA-Seq and scATAC-Seq with the SNARE-Seq data that we considered in Section 2.3 and consider the performance of LIGER and Signac, which performed comparatively well. We would note that LIGER and Signac have their own clustering strategies, separate from their dimensionality reduction, but we focus here on the results of their dimensionality reduction. We provided LIGER and Signac only the MOp data, as there is no clear way of including the SNARE-seq into LIGER and Signac without focusing on one of its modalities and adding extra batch correction steps. We compare the results to the clusters published with the scRNA-seq and scATAC-seq data in Yao et al. [2021] (Supplementary Figure 13, Additional File 1). We see that `Cobolt` generates a UMAP visualization that well represents rare subpopulations and respects broader cell classes. LIGER gives greater separation between cell types but splits several subtypes into far-away islands, such as for L5 PT, MGC, and Astro. Signac adopts an asymmetric strategy of transferring scRNA-seq labels to scATAC-seq data, and as a result, Signac performs well on major cell types but poorly on under-represented subpopulations in scRNA-seq such as astrocytes (Astro), which accounts for only 0.4% of the cells in scRNA-seq but 6.55% in scATAC-seq. Further, `Cobolt`, unlike LIGER and Signac, not only groups together the subtypes, but appears to also represent the three broader categories major GABAergic inhibitory neurons, glutamatergic excitatory neurons, and non-neurons.

These cluster identifications that we highlight from Yao et al. [2021] are relatively robust, well-known cell types, representing the large structural changes in the data, for which we expect most strategies to be able to detect reasonably well. On the other hand, our nearest-neighbor analysis emphasizes the performance at a high level of resolution. Putting both of these together points to the fact that `Cobolt` provides a superior integration of the datasets across a wide spectrum of resolutions.

## 3 Discussion

In this paper, we have shown that `Cobolt` successfully integrates multi-modal data and provides a representation that can be used for downstream analysis tasks, such as cell-type discovery. Pseudo-time estimation for reconstructing developmental order of cells Saelens et al. [2019], while not meaningful for the datasets we considered, is another important downstream application where the integrated representation of `Cobolt` allows the analysis of cells from different modalities. Future work could make use of the graphical model and inferred parameters to establish connections between features. For example, the probability vectors generated by *B*^(*i*)^ naturally provide a reduced-dimensional space of molecular features and can potentially help in the construction of gene networks.

We would note that while we have focused on the capability of `Cobolt` to analyze data from two-modalities, the underlying method can be extended to larger number of modalities and integration of different combinations of modalities, such as datasets with different pairs of modalities (see Methods, Section 5.1). Thus `Cobolt` provides a framework to integrate a wide range of varieties of multi-modality platforms as well as single-modality platforms.

`Cobolt` is available as a python package at https://github.com/epurdom/cobolt. All of the code used for the analysis is available as a github repository: https://github.com/epurdom/cobolt_manuscript.

## 4 Conclusions

We have shown that `Cobolt` is a flexible tool for analyzing multi-modality sequencing data, whether separately or integrated with single-modality data. `Cobolt` synthesizes the varied data into a single representation, preserving meaningful biological signal in the different modalities and at different resolutions. Moreover, this latent variable space is appropriate for standard downstream analysis techniques commonly used for analyzing cells without any further specialized adjustments, allowing `Cobolt` to fit into standard analysis pipelines.

## 5 Methods

While the most common application of joint-modality platforms consists of pairs of modalities (such as the example of mRNA-seq and ATAC-seq we described above), we will describe `Cobolt` in generality, assuming that there are *M* modalities.

### 5.1 Modeling Modality Dependency

For an individual cell *c*, we can (potentially) observe *M* vectors of data 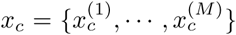, each vector of dimension *d*_1_, …, *d*_*M*_ corresponding to the number of features of each modality. We assume a Bayesian latent model, such that for each cell there is a latent variable *z*_*c*_ ∈ **R**^*K*^ representing the biological signal of the cell, where *z*_c_ is assumed drawn from a Gaussian prior distribution. Given *z*_c_, we assume that the data 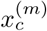 for each modality has an independent generative process, potentially different for each modality. Specifically, we assume that the data 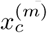 from each modality are conditionally independent given the common latent variable *z*_*c*_. That is,

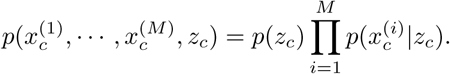

We use 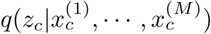 as a variational approximation of the posterior distribution 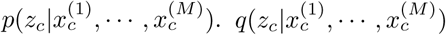, the encoder, is assumed Gaussian with parameters modeled as neural networks (i.e. Variational Autoencoder, VAE [Kingma and Welling, 2013]). This allows for estimation of the posterior distribution 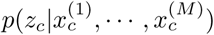 and the underlying latent variable for each cell *c*. The posterior mean of this distribution 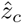 will be our summary of the shared representation across modalities.

Importantly, this model can be estimated even when not all of the input data contains all modalities. In this case, an individual cell *c* contains a subset of the modalities, S_*c*_ ⊂ {1, …, *M*}, and consists of data 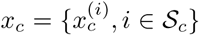. Without all modalities observed, the cell can contribute to the estimation of the model as its distribution can be explicitly written out:

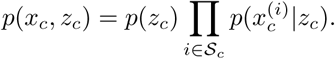

Further, we can estimate latent variables for such cells by using posterior distribution of *z*_*c*_ when conditioning only on the observed modalities, 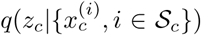. Instead of using separate neural networks for 2^*M*^ – 1 posterior distributions of different modality combinations, we adopt a technique introduced in Multimodal Variational Autoencoder (MVAE) [Wu and Goodman, 2018], which largely reduces the number of encoders to 2*M* (See Supplmentary Text for inference details).

As an example, if there are two modalities, mRNA-seq and ATAC-seq, and we have *n*_1_ cells with paired data from the joint modality platform, 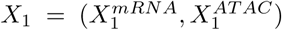; *n*_2_ cells with only mRNA measured, 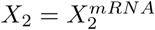; and *n*_3_ cells with only ATAC-seq measured, 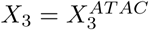. All *N* = *n*_1_ + *n*_2_ + *n*_3_ cells can be used in the estimation of the joint distribution of the latent variables, and estimates of the latent variables can be found as the mean of the relevant approximate posterior distributions:

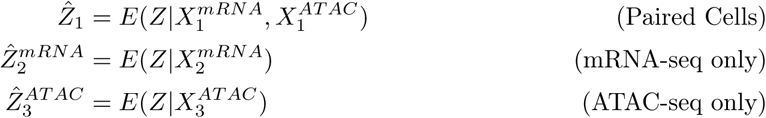

#### Correcting for missing modalities

In practice we find that the distributions 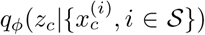 have subtle differences for different subsets S, i.e. the latent variables 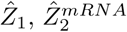, and 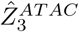 show distinct separations (Supplementary Figure 15, Additional File 1). One possibility could be due to platform differences between the different datasets that remain even after our batch correction (see Section 5.2). However, we also see differences in these distributions even if we only consider the joint-modality data, where we can estimate all of these posterior distributions on the same cells, 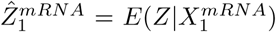 or 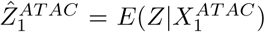 (Supplementary Figure 15, Additional File 1). Indeed there is nothing in the optimization of the posterior distribution that requires these different posterior distributions to be the same.

While the effects are small, these subtle differences can affect downstream analyses, e.g., in clustering cells for subtype discovery. Rather than directly force these posterior distributions to match in our estimation of the model, `Cobolt` fits the model as described above (using all of the data), then uses the paired data to train a prediction models that predict 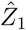 from the modality-specific estimates 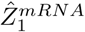 and 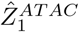. We then apply these prediction models to 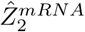 and 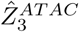 to obtain estimates 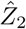 and 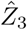 which are better aligned to be jointly analyzed in the same space. In practice, we find XGBoost [Chen and Guestrin, 2016] and k-nearest neighbors algorithm work equally well. We present results based on XGBoost (See Section 5.3 for details). We would note that there is little difference in performance when we predict coordinates into the ATAC-Seq space *E*(*Z*|*X*^*AT AC*^) or mRNA-Seq space *E*(*Z*|*X*^*mRNA*^), rather than the joint space 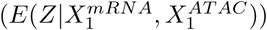, see Supplementary Figure 16 (Additional File 1).

### 5.2 Modeling Single Modality of Sparse Counts

The choice of the generative distribution *p*_*ψ*_(*x*^(*i*)^|*z*) should be chosen to reflect the data, and in principle can vary from modality to modality. For example, single-modality VAE models using zero-inflated negative binomial distributions (ZINB) [Lopez et al., 2018] have been proposed for scRNA-seq datasets to account for sparse count data. However, we found ZINB models performed less well for technologies that measure modalities like chromatin accessibility, which results in data with sparser counts and larger feature sizes than scRNA-seq. Therefore, we develop a latent model for these types of modalities inspired by the Latent Dirichlet Allocation (LDA) [Blei et al., 2003].

Our generative model for a single modality *i* starts by assuming that the counts measured on a cell are the mixture of the counts from different latent categories. In the genomic setting, these categories could correspond to biological processes of the cell, for example. Each category has a corresponding distribution of feature counts. The cumulative feature counts for a cell *c* are then the result of combining the counts across its categories, i.e. a mixture of the categories’ distributions. Specifically, each cell *c* has a latent probability vector *θ*_*c*_ ∈ [0, 1]^*K*^ describing the proportion of each category that makes up cell *c*. Each category *k* has a probability vector *σ*(*β*_*k*_) that provides the distribution of its feature counts. Here *σ* indicates the softmax function that transforms *β*_*k*_ to a probability vector that sums to 1. The observed vector of counts *x*_*c*_ are a multinomial draw with probabilities *π*_*c*_, where *π*_*c*_ = *σ*(*B*)*θ*_*c*_, and *B* = (*β*_1_, …, *β*_*K*_) is a matrix of the individual *β*_*k*_ vectors. To extend this model to multiple modalities, we assume a shared latent variable *z*_*c*_ that is common across modalities, but each modality has a different *B*^(*i*)^ that transforms the shared latent class probabilities into the feature space of the modality.

Furthermore, it is well known that there can be meaningful technical artifacts (“batch effects”) between different datasets on the same modality, for example due to differences between platforms or laboratory preparations. To counter this, our model also adjusts the sampling probabilities *σ*(*B*^(*i*)^)*θ*_*c*_ differently for data from different batches within the same modality *i*. We would note that the model can also take batch-corrected counts as input, such as are available for mRNA-Seq data, but we anticipate that for some modality types stand-alone batch correction techniques may not be as well developed. We evaluate the effect of our batch correction on the 10x Multiome data, which consists of two runs of 10x Multiome on the different patient input sampled at different times. This creates a batch effect between the multi-modality input and the single-modality input where we know the ground truth of how the single-modality data should be linked. We use the same nearest-neighbor analysis as in Figure 5B, with and without the batch correction terms and see much improved performance using the batch correction (Supplementary Figure 17, Additional File 1). This type of quantitative nearest-neighbor analysis is not possible for the SNARE-Seq data, since we do not have two different batches of paired multi-modality data, but we visually see large improvement due to the batch correction when analyzing the single-modality datasets jointly with the multimodality data (Supplementary Figure 18, Additional File 1).

The parameter *θ*_*c*_ is the latent variable describing the contributions of each category to cell *c*, and is shared across all modalities. In LDA models, it is typically assumed to have a Dirichlet prior distribution. However, we use a Laplace approximation to the Dirichlet introduced in ProdLDA [Srivastava and Sutton, 2017], which allows for incorporation into a VAE model. This prior assumes a latent variable *z*_*c*_ with a Gaussian prior, and sets *θ*_*c*_ = *σ*(*z*_*c*_), where *σ* is the softmax transformation. We use this approximation to the Dirichlet distribution to provide a multi-modality method appropriate for sparse sequence count data.

### 5.3 Data Processing and Method Implementation

#### SNARE-seq processing and annotation

We downloaded the processed counts of the adult mouse cerebral cortex data (10,309 cells) [Chen et al., 2019]. We applied quality filtering that retained cells having a number of genes detected greater than 20. For genes, we used the ones detected in more than 5 cells and have a total number of counts greater than 10. For peaks, we removed the ones having nonzero counts in more than 10% of cells or less than 5 cells. We performed clustering analysis using Seurat (version 3.2.2) on the gene expression modality. The data were normalized using `SCTransform` function with default parameters, followed by principal component analysis (PCA) using the default 3,000 variable features. Louvain algorithm was applied on the first 50 PCs with the resolution parameter equals 0.65. Cell type annotations are generated on the resulting 15 clusters using the marker genes [Yao et al., 2021]. We applied cisTopic (version 0.3.0) on the chromatin accessibility data with default parameters. Model selection was conducted based on log-likelihood using `runWrapLDAModels` and `selectModel` functions, and 30 topics are used in the results.

For the integration of SNARE-seq with the MOp data using `Cobolt`, we map the SNARE-seq counts to the peak set called on the MOp scATAC-seq data. Since peaks are typically called in a dataset-specific manner, the ideal integration strategy would be to redo the peak-calling with all datasets combined. However, in Supplementary Figure 10 we show that our simple alternative of mapping data to peaks called on a different dataset from the same system does not result in significant performance loss for `Cobolt` .

#### MOp Data Preprocessing

We downloaded the single-nucleus 10x v3 transcriptome dataset (90,266 cells) and the open chromatin dataset (15,731 cells, sample 171206_3C) [Yao et al., 2021]. For mRNA-seq quality control, we filtered cells that have less than 200 genes detected or have greater than 5% mitochondrial counts. For ATAC-seq, we utilized the `TSSEnrichment` and blacklist region reads calculation functionalities in Signac. We subsetted cells with the blacklist ratio less than 0.1, the number of unique molecular identifiers (UMIs) greater than 50, and the TSS enrichment score greater than 2 and less than 20. A total of 88,021 and 13,249 were retained for mRNA-seq and ATAC-seq, separately. To make annotation in the two datasets consistent, we merged the layer 2/3 IT and layer 4/5 IT subclusters in ATAC-seq data. For mRNA-seq, we merged Lamp5, Vip, and Sncg into one CGE cluster. When integrating the MOp datasets with the SNARE-seq data, we used only genes detected in both the scRNA-seq and the SNARE-seq datasets.

#### 10X PBMC data Preprocessing

PBMC datasets were downloaded from the 10X website (Multiome Chromium X, Multiome unsorted, scRNA-seq, scATAC-seq). For chromatin accessibility, we mapped the scATAC-seq reads and the Multiome unsorted reads to the peaks called on the Multiome Chromium X data. No quality filtering was applied to any of the four 10X datasets. Clusters with high mitochondrial expression are identified and annotated as low quality clusters, and clusters with the majority of cells identified by DoubletFinder McGinnis et al. [2019] are annotated as doublet clusters in the downstream clustering analysis.

#### Gene Activity Calculation

Gene activity matrix for chromatin accessibility is generated by counting the number of reads overlapping genes and their promoters using BEDOPS [Neph et al., 2012], where a promoter is defined as the region starting from the transcription start site (TSS) to 3,000 base pairs upstream of TSS.

#### Cobolt Network Architecture and Training

For each modality *i*, the encoder takes as input the log-plus-one transformed counts. We use one fully connected layer of size 128, followed by fully connected layers for mean 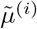 and log-variance 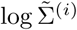. We tried networks with one or two hidden layers of varying sizes and found the results pretty stable. The decoders follow our probability model for sparse counts described in Section 5.2 (see also Equation 5, Supplementary Text, Additional File 2) and do not contain neural networks. We set the parameter of the Dirichlet prior to 50 divided by the number of latent variables *K*. The actual parameters used for the Gaussian prior are calculated using the Laplace approximation (see also Equation 3, Supplementary Text, Additional File 2). For the ELBO objective, we set the weighting terms *λ*^A^ reciprocal to the number of samples available for modality combination A. We set the hyperparameter weights *η* for conditional likelihood terms to 1. Adam optimizer is used, and we select a learning rate of 0.005 after tunning. We adopt a KL cost annealing schedule that linearly increases the weight of the KL term *γ* from 0 to 1 in the first 30 epochs. During training, we use a batch size of 128 and a fixed number of 100 epochs.

We note that the softmax transformation from *z*_*c*_ to *θ*_*c*_ is not a one-to-one transformation. Therefore, we scale 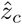 to mean 0 before the downstream correction, followed by clustering and visualization.

In correcting for missing modalities, we predict using XGBoost, setting the objective function to regression with squared loss, the learning rate to 0.8, and the maximum depth of a tree to 3. XGBoost is applied separately to each modality (scRNA-Seq and scATAC-Seq), resulting in our corrected estimates 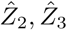.

The number of latent variables *K* was set to 10 in the SNARE-seq analysis (Section 2.2) and the method comparison analysis (Section 2.3.1) so as to be consistent with the default of several other methods for comparison purposes. We used *K* = 30 for the mouse cortex integration and *K* = 10 for the 10X PBMC integration (Section 2.3). Estimations of the data’s marginal likelihood were used to assist the selection of *K*.

For the mouse cortex data integration, we focused on genes and peaks that have top 30% average expression and removed the ones in the top 1%. For the SNARE-seq analysis and the 10X PBMC integration, all features were included in training. The choice of top features is less important here, which we found to have a small effect on the results.

#### Clustering and Visualization of Cobolt Results

Clusters were generated on the corrected latent variables using Louvain algorithm [Blondel et al., 2008]. We used the implementation of naive Louvain algorithm in `FindClusters` function from the R package Seurat. All parameters, other than the resolution controlling the number of clusters identified, were set to default. UMAPs were generated using the `umap` function from the R package uwot with the number of neighbors set to 30.

#### MOFA2, scMM, and BABEL analysis on the SNARE-seq data

Following the vignette of MOFA2, we used the top 2500 variable genes and cisTopic embeddings of the chromatin accessibility modality as input. Variable genes are selected using the `FindVariableFeatures` function from `Seurat` using selection method “vst”. The number of factors was set to 10. Two factors were identified as technical factors after inspecting their correlation with the total number of reads counts per cell. The UMAP representation was then generated using the rest of the factors with the number of neighbors set to 30. scMM was run with batch size equals 32, number of epochs equals 50, learning rate equals 10^−4^, number of latent dimensions equals 10, number of hidden dimension for gene expression equals 100, and number of hidden dimensions for chromatin accessibility equals 500. The parameters were chosen following the scMM paper. BABEL was run with the number of latent dimensions set to 10 and the batch size set to 256. Other parameters were chosen to follow the built-in SNARE-seq defaults.

#### Differential Analysis

DE analysis on gene expression and DA analysis on gene activities were performed using the Wilcoxon Rank Sum test followed by Bonferroni correction, implemented by the `FindMarkers` function in Seurat. Gene expression and gene activities were visualized by heatmaps using `DoHeatmap` in Seurat. DA analysis on the peaks for the SNARE-seq analysis was performed using Fisher’s exact test followed by the Benjamini–Hochberg procedure (following Chen et al. [2019]). Peak-by-cluster matrix was normalized by size factors calculated by Monocle Qiu et al. [2017] and visualized by heatmaps (following Chen et al. [2019]). Genes and peaks with adjusted *p*-values lower than 0.05 were called significant.

#### LIGER and Signac Analysis on the MOp Data

LIGER and Signac (Seurat) take as input gene-level count summaries from different modalities, such as gene expression or gene body methylation/chromatin accessibility measures. The input is different from `Cobolt`, which uses the peak summaries directly without needing to summarize at the gene level. Therefore, we applied these two methods on the gene expression and gene activity matrices, where the latter is defined as the summarized chromatin accessibility counts over gene and promoter regions.

We ran LIGER (version 0.5.0) using default parameters on the filtered data. Parameter K for factorization is set to 30 after inspecting the plot generated by function `suggestK`. Louvain clustering was performed by setting the resolution such that 17 clusters were obtained. We ran Signac (version 1.1.0) on the same filtered data. We first performed clustering analysis on the gene expression modality and then transferred the cluster labels to the open chromatin modality. Both the gene expression matrix and gene activity matrix were normalized by running `NormalizeData` followed by `ScaleData`. For the gene activity matrix, the `scale.factor` parameter was set to the median of UMI distribution as suggested by the vignette. Other parameters were kept as default. We then ran `FindTransferAnchors` with the default number of dimensions equals 30, which is the same as used for LIGER. Finally, we ran `TransferData` with weight reduction set to “cca” or the Latent Semantic Indexing (LSI) from analyzing the peak matrix. Results using LSI are presented in the paper as it performed better than the “cca” option.

#### Test training split for method comparision

For scMM and BABEL, which do not allow the single-modality data to be used in the training of the model, we assign 20% of the cells as paired modality data that is used for the training set; the trained model was then used to generate the embedding on the rest of the cells without providing the pairing information. This provided separate estimates for the unpaired mRNA and ATAC modalities, respectively, with which we evaluated whether the paired cells were close together. For `Cobolt`, which allows the use of single-modality data for the training of the model, we assign 20% of the cells as paired modality data and the other 80% were given as single-modality data, and then we trained the model on all of the cells. For LIGER and Signac, which are not designed for multiple-modality data, we hide the pairing information on all cells, and treat all of the cells as if they were collected on different cells. To be comparable with the other methods, we only evaluated their performance on the 80% of cells treated by the other methods as single-modality data.

## Acknowledgements

The authors would like to thank Jean-Philippe Vert for useful early discussions on the analysis of multimodality data, and Hector Roux de Bezieux for assistance in accessing the MOp data.

## Availability of data and materials

An open-source package `Cobolt`, implemented in Python, is available on GitHub (https://github.com/epurdom/cobolt). Code scripts for reproducing results presented in the paper are publicly accessible on GitHub (https://github.com/epurdom/cobolt_manuscript).

Datasets used in this paper are all publicly available. The SNARE-seq data can be downloaded from Gene Expression Omnibus with accession number GSE126074. The MOp scRNA-seq and scATAC-seq data can be downloaded from NeMO Archive with accession number nemo:dat-ch1nqb7. The 10X datasets are available on the 10x Genomics website (Multiome Chromium X, Multiome unsorted, scRNA-seq, scATAC-seq).

## Ethics approval and consent to participate

Not applicable.

## Consent for publication

Not applicable.

## Authors’ contributions

EP formulated the problem and supervised the work. BG developed the model and performed the analysis, with inputs from all authors. BG and YZ developed the model inference and coded the initial implementation. BG implemented the package. BG and EP drafted and revised the manuscript. YZ revised the manuscript and figures. All authors read and approved the final manuscript.

## Additional Files

**Additional file 1 — Supplementary Figures**

PDF file containing Supplementary Figures referenced in the main text.

**Additional file 2 — Supplementary Text**

PDF file containing additional details regarding the `Cobolt` algorithm.

## References

[1] 10x Genomics. Pbmcs from human (atac v1.1, chromium x), single cell atac dataset by cell ranger atac 2.0.0. 2021, August 9a.

[2] 10x Genomics. Pbmcs from human (3’ ht v3.1, chromium x), single cell gene expression dataset by cell ranger 6.1.0. 2021, August 9b.

[3] 10x Genomics. Pbmcs from human (multiome v1.0, chromium x), single cell multiome atac + gene expression dataset by cell ranger arc 2.0.0. 2021, August 9c.

[4] 10x Genomics. Pbmcs from human (no cell sorting, chromium next gem), single cell multiome atac + gene expression dataset by cell ranger arc 2.0.0. 2021, May 3.

[5] Ricard Argelaguet, Damien Arnol, Danila Bredikhin, Yonatan Deloro, Britta Velten, John C Marioni, and Oliver Stegle. Mofa+: a statistical framework for comprehensive integration of multi-modal single-cell data. Genome Biology, 21:1–17, 2020.

[6] Daniel Backenroth, Zihuai He, Krzysztof Kiryluk, Valentina Boeva, Lynn Pethukova, Ekta Khurana, Angela Christiano, Joseph D Buxbaum, and Iuliana Ionita-Laza. Fun-lda: a latent dirichlet allocation model for predicting tissue-specific functional effects of noncoding variation: methods and applications. The American Journal of Human Genetics, 102(5):920–942, 2018.

[7] Arielle Baker, Brian Kalmbach, Mieko Morishima, Juhyun Kim, Ashley Juavinett, Nuo Li, and Nikolai Dembrow. Specialized subpopulations of deep-layer pyramidal neurons in the neocortex: Bridging cellular properties to functional consequences. Journal of Neuroscience, 38(24):5441–5455, 2018. ISSN 15292401. doi: 10.1523/JNEUROSCI.0150-18.2018.

[8] T Grant Belgard, Ana C Marques, Peter L Oliver, Hatice Ozel Abaan, Tamara M Sirey, Anna Hoerder-Suabedissen, Fernando García-Moreno, Zoltán Molnár, Elliott H Margulies, and Chris P Ponting. A transcriptomic atlas of mouse neocortical layers. Neuron, 71(4):605–616, 08 2011. doi: 10.1016/j.neuron.2011.06.039. URL https://pubmed.ncbi.nlm.nih.gov/21867878.

[9] David M Blei, Andrew Y Ng, and Michael I Jordan. Latent dirichlet allocation. Journal of machine Learning research, 3(Jan):993–1022, 2003.

[10] Vincent D Blondel, Jean-Loup Guillaume, Renaud Lambiotte, and Etienne Lefebvre. Fast unfolding of communities in large networks. Journal of statistical mechanics: theory and experiment, 2008(10):P10008, 2008.

[11] Junyue Cao, Darren A Cusanovich, Vijay Ramani, Delasa Aghamirzaie, Hannah A Pliner, Andrew J Hill, Riza M Daza, Jose L McFaline-Figueroa, Jonathan S Packer, Lena Christiansen, et al. Joint profiling of chromatin accessibility and gene expression in thousands of single cells. Science, 361(6409):1380–1385, 2018.

[12] Song Chen, Blue B Lake, and Kun Zhang. High-throughput sequencing of the transcriptome and chromatin accessibility in the same cell. Nature biotechnology, 37(12):1452–1457, 2019.

[13] Tianqi Chen and Carlos Guestrin. Xgboost: A scalable tree boosting system. In Proceedings of the 22nd acm sigkdd international conference on knowledge discovery and data mining, pages 785–794, 2016.

[14] Stephen J Clark, Ricard Argelaguet, Chantriolnt-Andreas Kapourani, Thomas M Stubbs, Heather J Lee, Celia Alda-Catalinas, Felix Krueger, Guido Sanguinetti, Gavin Kelsey, John C Marioni, et al. scnmt-seqenables joint profiling of chromatin accessibility dna methylation and transcription in single cells. Nature communications, 9(1):1–9, 2018.

[15] Jiarui Ding, Xian Adiconis, Sean K Simmons, Monika S Kowalczyk, Cynthia C Hession, Nemanja D Marjanovic, Travis K Hughes, Marc H Wadsworth, Tyler Burks, Lan T Nguyen, et al. Systematic comparison of single-cell and single-nucleus rna-sequencing methods. Nature biotechnology, 38(6):737–746, 2020.

[16] Siavash Fazel Darbandi, Sarah E. Robinson Schwartz, Qihao Qi, Rinaldo Catta-Preta, Emily Ling Lin Pai, Jeffrey D. Mandell, Amanda Everitt, Anna Rubin, Rebecca A. Krasnoff, Sol Katzman, David Tastad, Alex S. Nord, A. Jeremy Willsey, Bin Chen, Matthew W. State, Vikaas S. Sohal, and John L.R. Rubenstein. Neonatal Tbr1 Dosage Controls Cortical Layer 6 Connectivity. Neuron, 100(4):831–845.e7, 2018. ISSN 10974199. doi: 10.1016/j.neuron.2018.09.027. URL https://doi.org/10.1016/j.neuron.2018.09.027.

[17] Oscar Franzén, Li-Ming Gan, and Johan LM Björkegren. Panglaodb: a web server for exploration of mouse and human single-cell rna sequencing data. Database, 2019, 2019.

[18] Adam Gayoso, Zoë Steier, Romain Lopez, Jeffrey Regier, Kristopher L Nazor, Aaron Streets, and Nir Yosef. Joint probabilistic modeling of single-cell multi-omic data with totalvi. Nature Methods, pages 1–11, 2021.

[19] Carmen Bravo González-Blas, Liesbeth Minnoye, Dafni Papasokrati, Sara Aibar, Gert Hulselmans, Valerie Christiaens, Kristofer Davie, Jasper Wouters, and Stein Aerts. cistopic: cis-regulatory topic modeling on single-cell atac-seq data. Nature methods, 16(5):397–400, 2019.

[20] Yuhan Hao, Stephanie Hao, Erica Andersen-Nissen, William M Mauck, Shiwei Zheng, Andrew Butler, Maddie Jane Lee, Aaron J Wilk, Charlotte Darby, Michael Zagar, et al. Integrated analysis of multimodal single-cell data. bioRxiv, 2020.

[21] Diederik P Kingma and Max Welling. Auto-encoding variational bayes. arXiv preprint 1312.6114, 2013.

[22] Jeongwoo Lee, Daehee Hwang, et al. Single-cell multiomics: technologies and data analysis methods. Experimental & Molecular Medicine, 52(9):1428–1442, 2020.

[23] Romain Lopez, Jeffrey Regier, Michael B Cole, Michael I Jordan, and Nir Yosef. Deep generative modeling for single-cell transcriptomics. Nature methods, 15(12):1053–1058, 2018.

[24] Christopher S McGinnis, Lyndsay M Murrow, and Zev J Gartner. Doubletfinder: doublet detection in single-cell rna sequencing data using artificial nearest neighbors. Cell systems, 8(4):329–337, 2019.

[25] Leland McInnes, John Healy, and James Melville. Umap: Uniform manifold approximation and projection for dimension reduction. arXiv preprint 1802.03426, 2018.

[26] Kodai Minoura, Ko Abe, Hyunha Nam, Hiroyoshi Nishikawa, and Teppei Shimamura. Scmm: Mixture-of-experts multimodal deep generative model for single-cell multiomics data analysis. Available at SSRN 3806072, 2021.

[27] Shane Neph, M Scott Kuehn, Alex P Reynolds, Eric Haugen, Robert E Thurman, Audra K Johnson, Eric Rynes, Matthew T Maurano, Jeff Vierstra, Sean Thomas, et al. Bedops: high-performance genomic feature operations. Bioinformatics, 28(14):1919–1920, 2012.

[28] Hannah A Pliner, Jay Shendure, and Cole Trapnell. Supervised classification enables rapid annotation of cell atlases. Nature methods, 16(10):983–986, 2019.

[29] Xiaojie Qiu, Andrew Hill, Jonathan Packer, Dejun Lin, Yi-An Ma, and Cole Trapnell. Single-cell mrna quantification and differential analysis with census. Nature methods, 14(3):309–315, 2017.

[30] Wouter Saelens, Robrecht Cannoodt, Helena Todorov, and Yvan Saeys. A comparison of single-cell trajectory inference methods. Nature Biotechnology, 37(5):547–554, 2019. doi: 10.1038/s41587-019-0071-9. URL https://doi.org/10.1038/s41587-019-0071-9.

[31] Staci A. Sorensen, Amy Bernard, Vilas Menon, Joshua J. Royall, Katie J. Glattfelder, Tsega Desta, Karla Hirokawa, Marty Mortrud, Jeremy A. Miller, Hongkui Zeng, John G. Hohmann, Allan R. Jones, and Ed S. Lein. Correlated gene expression and target specificity demonstrate excitatory projection neuron diversity. Cerebral Cortex, 25(2):433–449, 2015. ISSN 14602199. doi: 10.1093/cercor/bht243.

[32] Akash Srivastava and Charles Sutton. Autoencoding variational inference for topic models. International Conference on Learning Representations, 2017.

[33] Marlon Stoeckius, Christoph Hafemeister, William Stephenson, Brian Houck-Loomis, Pratip K Chattopadhyay, Harold Swerdlow, Rahul Satija, and Peter Smibert. Simultaneous epitope and transcriptome measurement in single cells. Nature methods, 14(9):865–868, 2017.

[34] Tim Stuart, Andrew Butler, Paul Hoffman, Christoph Hafemeister, Efthymia Papalexi, William M Mauck III, Yuhan Hao, Marlon Stoeckius, Peter Smibert, and Rahul Satija. Comprehensive integration of single-cell data. Cell, 177(7):1888–1902, 2019.

[35] Tim Stuart, Avi Srivastava, Caleb Lareau, and Rahul Satija. Multimodal single-cell chromatin analysis with signac. bioRxiv, 2020.

[36] Zhe Sun, Ting Wang, Ke Deng, Xiao-Feng Wang, Robert Lafyatis, Ying Ding, Ming Hu, and Wei Chen. Dimm-sc: a dirichlet mixture model for clustering droplet-based single cell transcriptomic data. Bioinformatics, 34(1):139–146, 2018.

[37] Bosiljka Tasic, Vilas Menon, Thuc Nghi Nguyen, Tae Kyung Kim, Tim Jarsky, Zizhen Yao, Boaz Levi, Lucas T Gray, Staci A Sorensen, Tim Dolbeare, et al. Adult mouse cortical cell taxonomy revealed by single cell transcriptomics. Nature neuroscience, 19(2):335–346, 2016.

[38] Bosiljka Tasic, Zizhen Yao, Lucas T. Graybuck, Kimberly A. Smith, Thuc Nghi Nguyen, Darren Bertagnolli, Jeff Goldy, Emma Garren, Michael N. Economo, Sarada Viswanathan, Osnat Penn, Trygve Bakken, Vilas Menon, Jeremy Miller, Olivia Fong, Karla E. Hirokawa, Kanan Lathia, Christine Rimorin, Michael Tieu, Rachael Larsen, Tamara Casper, Eliza Barkan, Matthew Kroll, Sheana Parry, Nadiya V. Shapovalova, Daniel Hirschstein, Julie Pendergraft, Heather A. Sullivan, Tae Kyung Kim, Aaron Szafer, Nick Dee, Peter Groblewski, Ian Wickersham, Ali Cetin, Julie A. Harris, Boaz P. Levi, Susan M. Sunkin, Linda Madisen, Tanya L. Daigle, Loren Looger, Amy Bernard, John Phillips, Ed Lein, Michael Hawrylycz, Karel Svoboda, Allan R. Jones, Christof Koch, and Hongkui Zeng. Shared and distinct transcriptomic cell types across neocortical areas. Nature, 563(7729):72–78, 2018. ISSN 14764687. doi: 10.1038/s41586-018-0654-5. URL http://dx.doi.org/10.1038/s41586-018-0654-5.

[39] Xinjun Wang, Zhe Sun, Yanfu Zhang, Zhongli Xu, Hongyi Xin, Heng Huang, Richard H Duerr, Kong Chen, Ying Ding, and Wei Chen. Brem-sc: a bayesian random effects mixture model for joint clustering single cell multi-omics data. Nucleic acids research, 48(11):5814–5824, 2020.

[40] Joshua D Welch, Velina Kozareva, Ashley Ferreira, Charles Vanderburg, Carly Martin, and Evan Z Macosko. Single-cell multi-omic integration compares and contrasts features of brain cell identity. Cell, 177(7):1873– 1887, 2019.

[41] Kevin E Wu, Kathryn E Yost, Howard Y Chang, and James Zou. Babel enables cross-modality translation between multiomic profiles at single-cell resolution. Proceedings of the National Academy of Sciences, 118 (15), 2021.

[42] Mike Wu and Noah Goodman. Multimodal generative models for scalable weakly-supervised learning. In Advances in Neural Information Processing Systems, pages 5575–5585, 2018.

[43] Zizhen Yao, Hanqing Liu, Fangming Xie, Stephan Fischer, Ricky S. Adkins, Andrew I. Aldridge, Seth A. Ament, Anna Bartlett, M. Margarita Behrens, Koen Van den Berge, Darren Bertagnolli, Hector Roux de Bézieux, Tommaso Biancalani, A. Sina Booeshaghi, Héctor Corrada Bravo, Tamara Casper, Carlo Colantuoni, Jonathan Crabtree, Heather Creasy, Kirsten Crichton, Megan Crow, Nick Dee, Elizabeth L. Dougherty, Wayne I. Doyle, Sandrine Dudoit, Rongxin Fang, Victor Felix, Olivia Fong, Michelle Giglio, Jeff Goldy, Mike Hawrylycz, Brian R. Herb, Ronna Hertzano, Xiaomeng Hou, Qiwen Hu, Jayaram Kancherla, Matthew Kroll, Kanan Lathia, Yang Eric Li, Jacinta D. Lucero, Chongyuan Luo, Anup Mahurkar, Delissa McMillen, Naeem M. Nadaf, Joseph R. Nery, Thuc Nghi Nguyen, Sheng-Yong Niu, Vasilis Ntranos, Joshua Orvis, Julia K. Osteen, Thanh Pham, Antonio Pinto-Duarte, Olivier Poirion, Sebastian Preissl, Elizabeth Purdom, Christine Rimorin, Davide Risso, Angeline C. Rivkin, Kimberly Smith, Kelly Street, Josef Sulc, Valentine Svensson, Michael Tieu, Amy Torkelson, Herman Tung, Eeshit Dhaval Vaishnav, Charles R. Vanderburg, Cindy van Velthoven, Xinxin Wang, Owen R. White, Z. Josh Huang, Peter V. Kharchenko, Lior Pachter, John Ngai, Aviv Regev, Bosiljka Tasic, Joshua D. Welch, Jesse Gillis, Evan Z. Macosko, Bing Ren, Joseph R. Ecker, Hongkui Zeng, and Eran A. Mukamel. A transcriptomic and epigenomic cell atlas of the mouse primary motor cortex. Nature, 598(7879):103–110, 2021. ISSN 0028-0836. doi: 10.1038/s41586-021-03500-8. URL http://dx.doi.org/10.1038/s41586-021-03500-8.

[44] Sohiya Yotsukura, Seitaro Nomura, Hiroyuki Aburatani, Koji Tsuda, et al. Celltree: an r/bioconductor package to infer the hierarchical structure of cell populations from single-cell rna-seq data. BMC bioinformatics, 17(1):363, 2016.

[45] Hongkui Zeng, Elaine H. Shen, John G. Hohmann, Seung Wook Oh, Amy Bernard, Joshua J. Royall, Katie J. Glattfelder, Susan M. Sunkin, John A. Morris, Angela L. Guillozet-Bongaarts, Kimberly A. Smith, Amanda J. Ebbert, Beryl Swanson, Leonard Kuan, Damon T. Page, Caroline C. Overly, Ed S. Lein, Michael J. Hawrylycz, Patrick R. Hof, Thomas M. Hyde, Joel E. Kleinman, and Allan R. Jones. Large-scale cellular-resolution gene profiling in human neocortex reveals species-specific molecular signatures. Cell, 149(2):483–496, 2021/10/19 2012. doi: 10.1016/j.cell.2012.02.052. URL https://doi.org/10.1016/j.cell.2012.02.052.

[46] Chenxu Zhu, Miao Yu, Hui Huang, Ivan Juric, Armen Abnousi, Rong Hu, Jacinta Lucero, M Margarita Behrens, Ming Hu, and Bing Ren. An ultra high-throughput method for single-cell joint analysis of open chromatin and transcriptome. Nature structural & molecular biology, 26(11):1063–1070, 2019.

[47] Chunman Zuo and Luonan Chen. Deep-joint-learning analysis model of single cell transcriptome and open chromatin accessibility data. Briefings in Bioinformatics, 22(4):bbaa287, 2021.

