## Supplementary Figures for "Cobolt: Joint analysis of multimodal single-cell sequencing data"

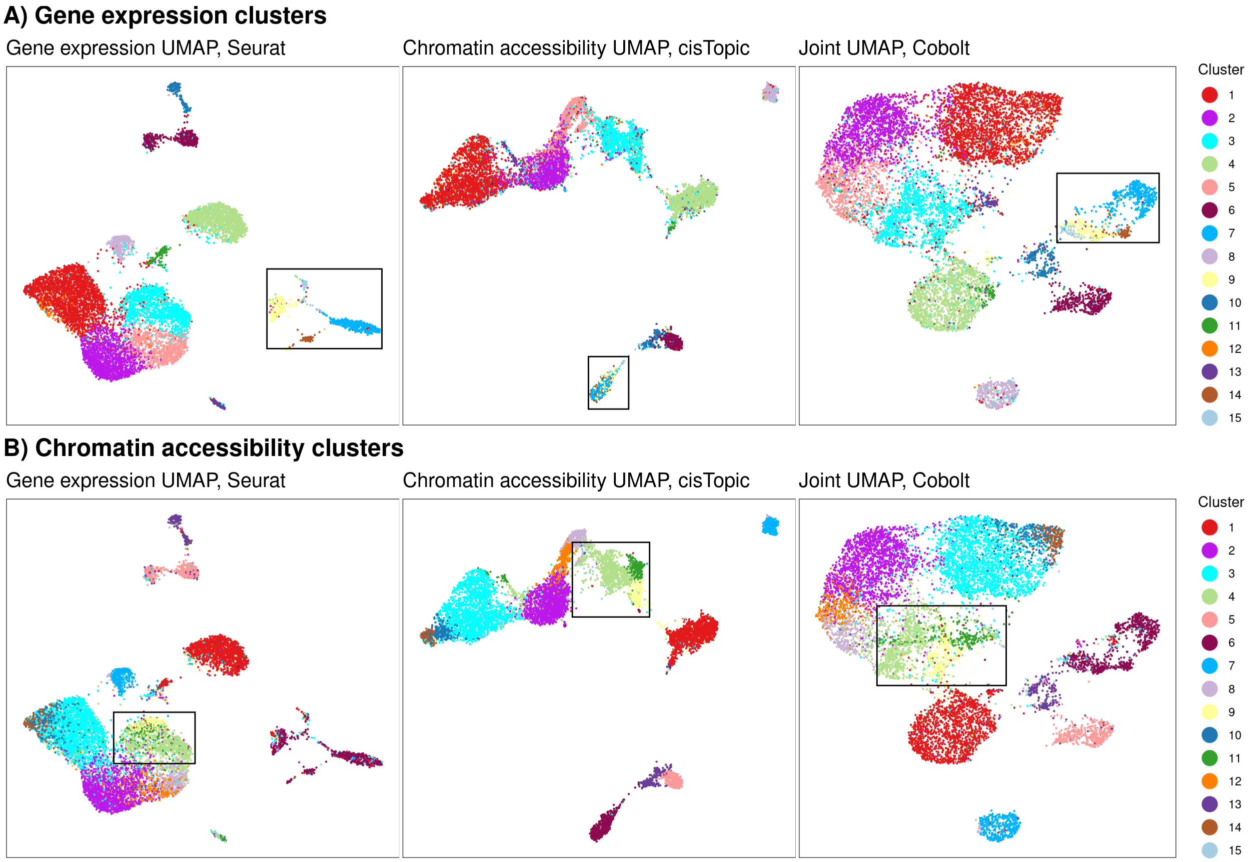


Supplementary Figure 1: Plotted are UMAP visualizations of the reduced dimensionality space created by analyzing only gene expression data using Seurat [Stuart et al., 2019b] and only chromatin accessibility using cisTopic [González-Blas et al., 2019], and jointly using both modalities (`Cobolt`). The cells are color-coded by the cluster they are assigned to based on clustering of A) gene expression modality and B) chromatin accessibility. We note that the cluster colors are randomly and separately assigned for panel A and B. In both cases, we see distinct clusters that are found in one modality but are not reflected in the other modality. Highlighted in the panels of A) are cells that show separated clusters in the gene expression, but not in chromatin accessibility; these cells show gene markers that show that these clusters correspond to the non-neuronal celltypes including astrocytes (Astro), microglial cells (MGC), oligodendrocyte precursors (OPC), and oligodendrocytes (Oligo) (See Supplementary Figure 2A). Highlighted in B) are cells that are well separated in chromatin accessibility but not in gene expression, and are potentially a subtype of Layer 6 cells (see Supplementary Figure 2B,C). Both sets of highlighted cells show separation in the joint analysis of `Cobolt`. In both these cases, the results of the `Cobolt` analysis shows that `Cobolt` integrates both modalities to find these modality-specific subtype differences: the two highlighted cell populations mentioned above are both distinguishable in the `Cobolt` reduced dimensions, and their cluster median silhouette widths are improved compared to single modality analysis (Supplementary Figure 3). These results demonstrate the power of a joint analysis of modalities using `Cobolt` for a comprehensive understanding of subtypes.

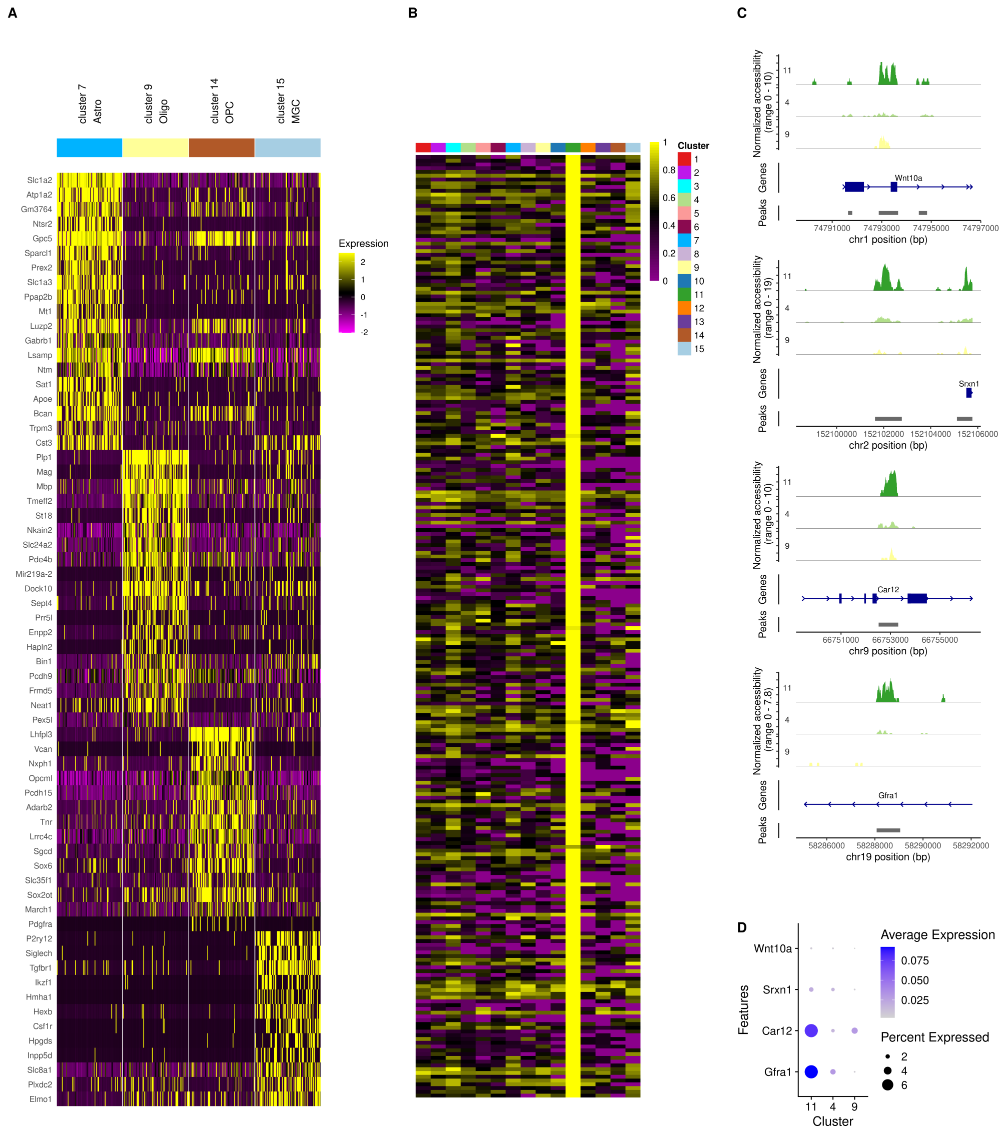

Supplementary Figure 2: **Clusters of SNARE-seq found by `Cobolt` and not found in one of the single modality analyses**. (A) Top genes from a differential expression analysis performed on clusters mentioned in the text as correctly identified by `Cobolt`. The cluster colors correspond to the mRNA clusters colored in Supplementary Figure 1A found by clustering the mRNA expression only using Seurat (i.e. not using the joint representation of `Cobolt`). The clustering of only the mRNA expression from the SNARE-seq data distinguishes clusters 7, 9, 14, and 15 (blue, yellow, brown and light blue) which are not apparent in the chromatin accessibility data, but are distinguished by `Cobolt`. *Atp1a2, Bcan*, and *Cst3* are highly expressed in cluster 7 (blue) and are markers of astrocytes (Astro) [Zeng et al., 2012]; *Mbp* is highly expressed in cluster 9 (yellow) and a marker of oligodendrocytes (Oligo); *Pdgfra* is highly expressed in cluster 14 (brown) and is a marker of oligodendrocyte precursors (OPC) [Yao et al., 2021]; and *Siglech* and *P2ry12* are highly expressed in cluster 15 (light blue) and are markers of microglial cells (MGC) [Tasic et al., 2018]. (B, C, D) Clustering of only the chromatin accessibility from the SNARE-seq data (Supplementary Figure 1B) discovers cluster 11 (green) which is not apparent in the mRNA expression clusters, but is found by `Cobolt`. Plotted in (B) are normalized accessibility levels of the top 200 DA peaks between cluster 11 (dark green) and its nearby cluster 4 and 9. Cluster 4, 9, and 11 are all identified as Layer 5/6 cells from their gene expression profile. We show examples of top peaks that are close to genes in (C) and the corresponding gene expression levels in (D), where *Car12* is a marker gene of Layer 6 subtype L6 Car12 Tasic et al. [2016].

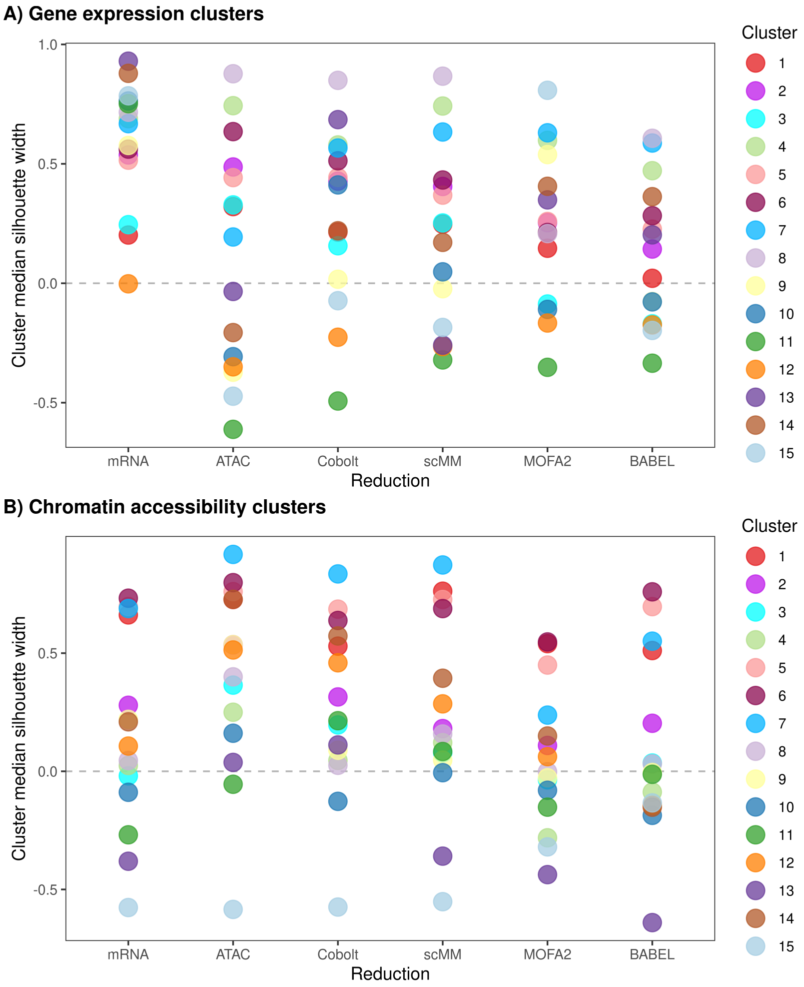


Supplementary Figure 3: The median silhouette width of SNARE-seq clusters generated using A) gene expression modality and B) chromatin accessibility. For each of the clustering, we calculate the median silhouette width per cluster on reduced dimensionality space created by analyzing only gene expression data (using Seurat), only chromatin accessibility (using cisTopic), and jointly with both modalities using `Cobolt`, scMM, MOFA2, and BABEL. Cluster colors are assigned the same as in Supplementary Figure 1.

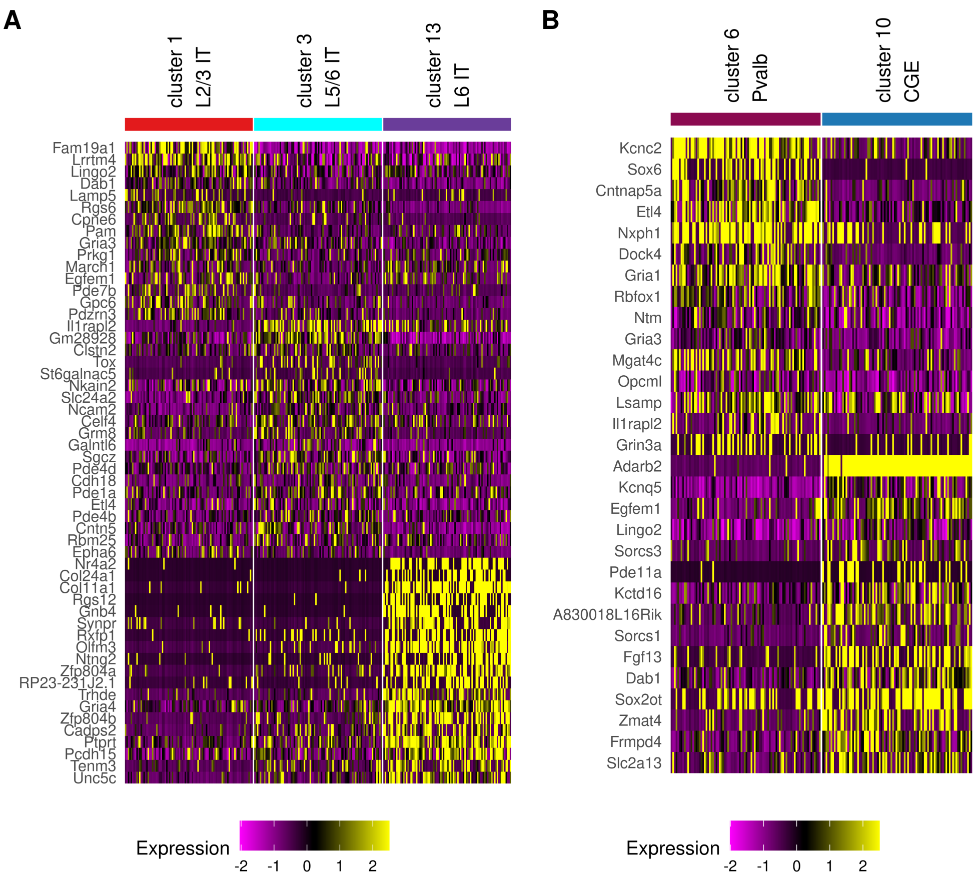


Supplementary Figure 4: **Clusters of SNARE-seq found by `Cobolt` and not found by other multi-modality methods**. Each heatmap shows top genes from a differential expression analysis performed on clusters mentioned in the text as correctly identified by `Cobolt` but missed by another multi-modality method. The cluster colors correspond to the mRNA clusters colored in Figure 2A and Supplementary Figure 1A which were found by clustering only the mRNA expression using Seurat (i.e. not using the joint representation of `Cobolt`). (A) `Cobolt` distinguishes the cells in cluster 13 (purple) from cells in the surrounding clusters 1 (red) and 3 (cyan) which scMM does not distinguish. Cluster 13 shows strong expression of *Col24a1, Gnb4, Rxfp1, Nr4a2, Ntng2, Car3* which are known markers of Layer 6 cells (B) `Cobolt` distinguishes the cells in cluster 5 (mauve) from cells in cluster 9 (blue) which MOFA2 does not distinguish. Cluster 5 shows expression of Sox6, a marker of Pvalb cells while cluster 9 shows expression of Adarb2, a marker of CGE cells [Yao et al., 2021].

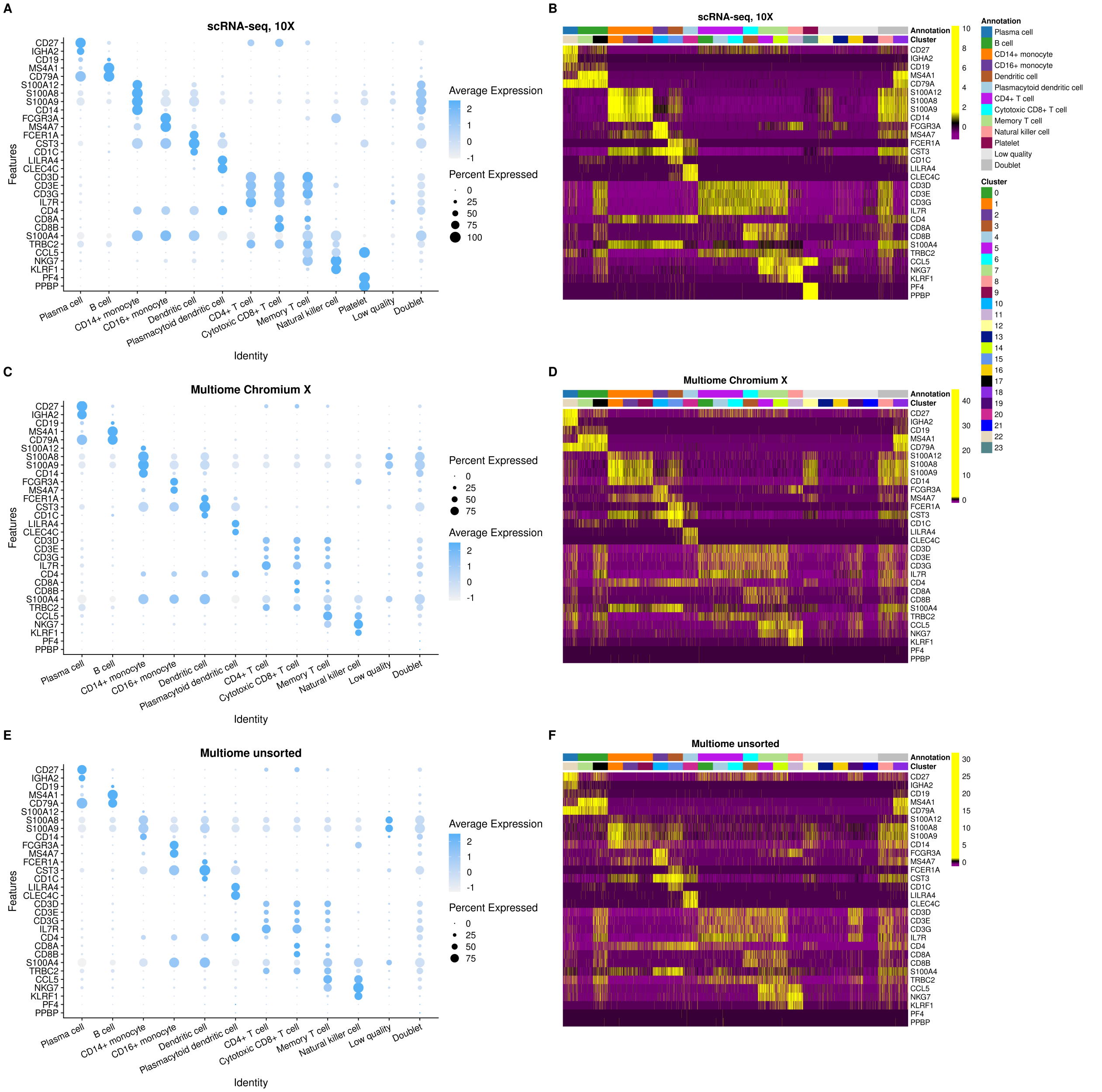


Supplementary Figure 5: **Expression of known marker genes of PMBCs, mRNA Expression** These plots show the mRNA expression of known marker genes of common cell types expected in PMBCs [Ding et al., 2020, Stuart et al., 2019a, Pliner et al., 2019, Franzén et al., 2019] for cells from the three PBMC mRNA expression datasets: (A-B) scRNA-seq, (C-D) Multiome Chromium X and (E-F) Multiome unsorted. The heatmaps (right) show the expression of cells for these gene markers in the clusters identified by the clustering of the `Cobolt` integration of the PBMC datasets. The results of clustering the `Cobolt` integration of the PBMC datasets are indicated above the cells; also indicated above are our assignment of these clusters to the known cell types based on these markers. The dot plots (left) show the expression of these gene markers for cells based on their assignment of to these known celltypes, illustrating that the assignments of clusters to known cell types result in cell types that follow known patterns in these gene markers.

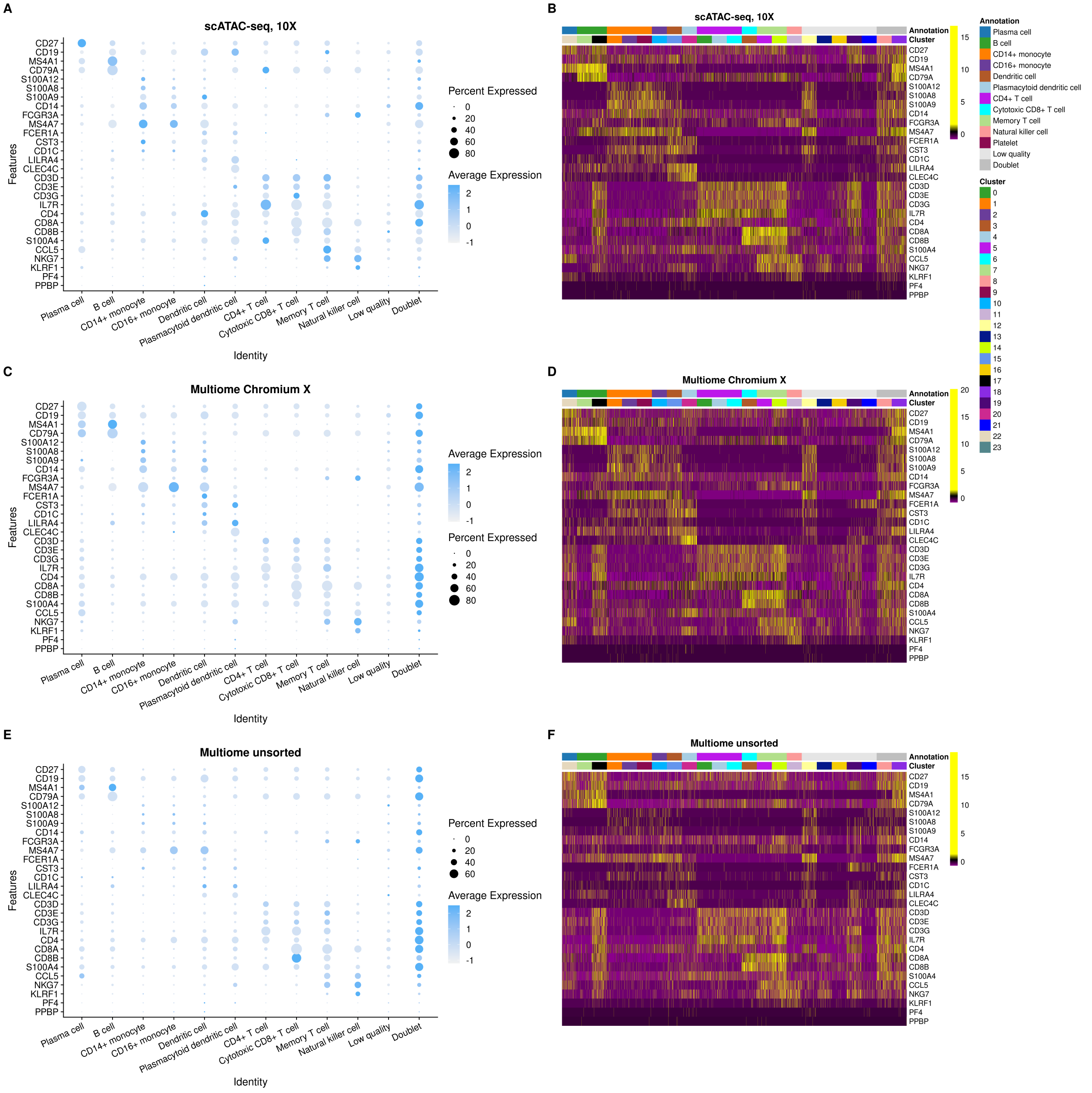


Supplementary Figure 6: **Expression of known marker genes of PMBCs, Chromatin Accessibility** These plots show the chromatin accessibility of known marker genes of common cell types expected in PMBCs [Ding et al., 2020, Stuart et al., 2019a, Pliner et al., 2019, Franzén et al., 2019] for cells from the three PBMC chromatim accessibility datasets: (A-B) scATAC-seq, (C-D) Multiome Chromium X and (E-F) Multiome unsorted. Chromatin accessibility levels are summarized over gene body and promoter regions. See Supplementary Figure 5 for futher details.

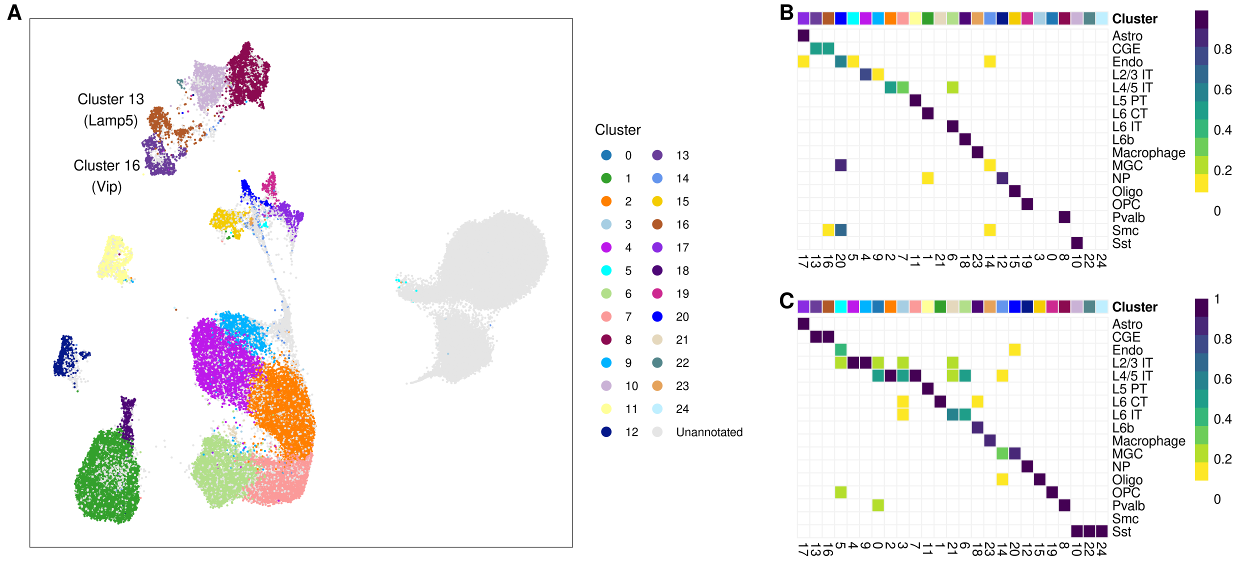


Supplementary Figure 7: `Cobolt` **clustering results on the mouse cortex data integration with cluster 13 and cluster 16 discussed in Figure 4 highlighted.** A) UMAP visualization colored by clusters. B, C) Concordance matrices between clustering results and annotations, scaled to sum 1 by (B) rows and (C) columns.

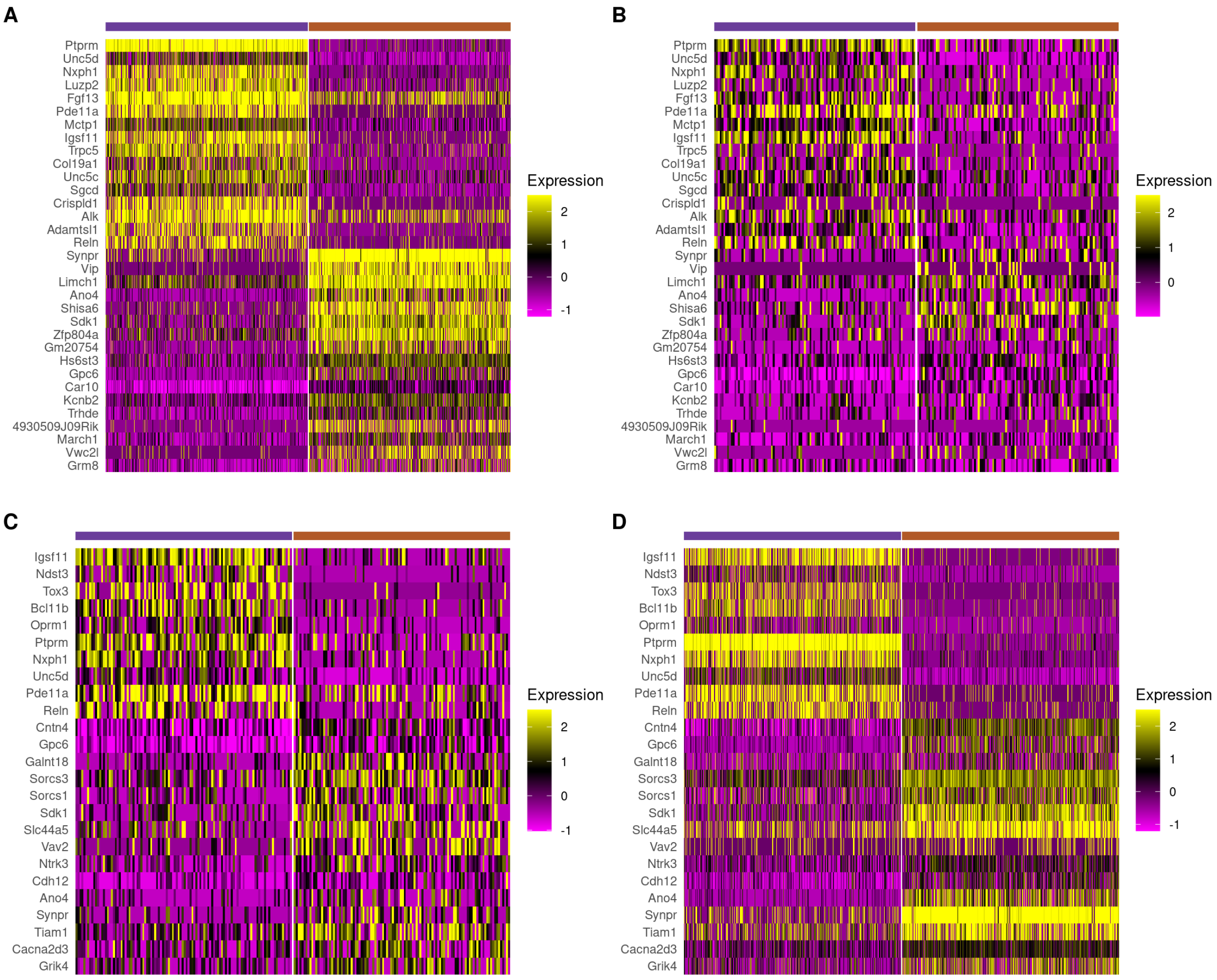


Supplementary Figure 8: **The DE and DA results between the *Vip* (cluster 16, brown) and the *Lamp5* (cluster 13, purple) clusters from the mouse cortex analysis.** A) The mRNA expression of the top DE genes between clusters 13 and 16. B) The ATAC chromatin accessibility levels of the genes identified in the DE analysis (shown in A)). C) The levels of ATAC chromatin accessibility of the top DA genes between clusters 13 and 16. D) The mRNA expression of the genes identified in the DA analysis (shown in C)). We note that all the top genes have the same direction of fold changes in the scRNA-seq and the scATAC-seq data, i.e., the genes with lower/higher gene expression in cluster 13 compared to cluster 16 are also less/more accessible.

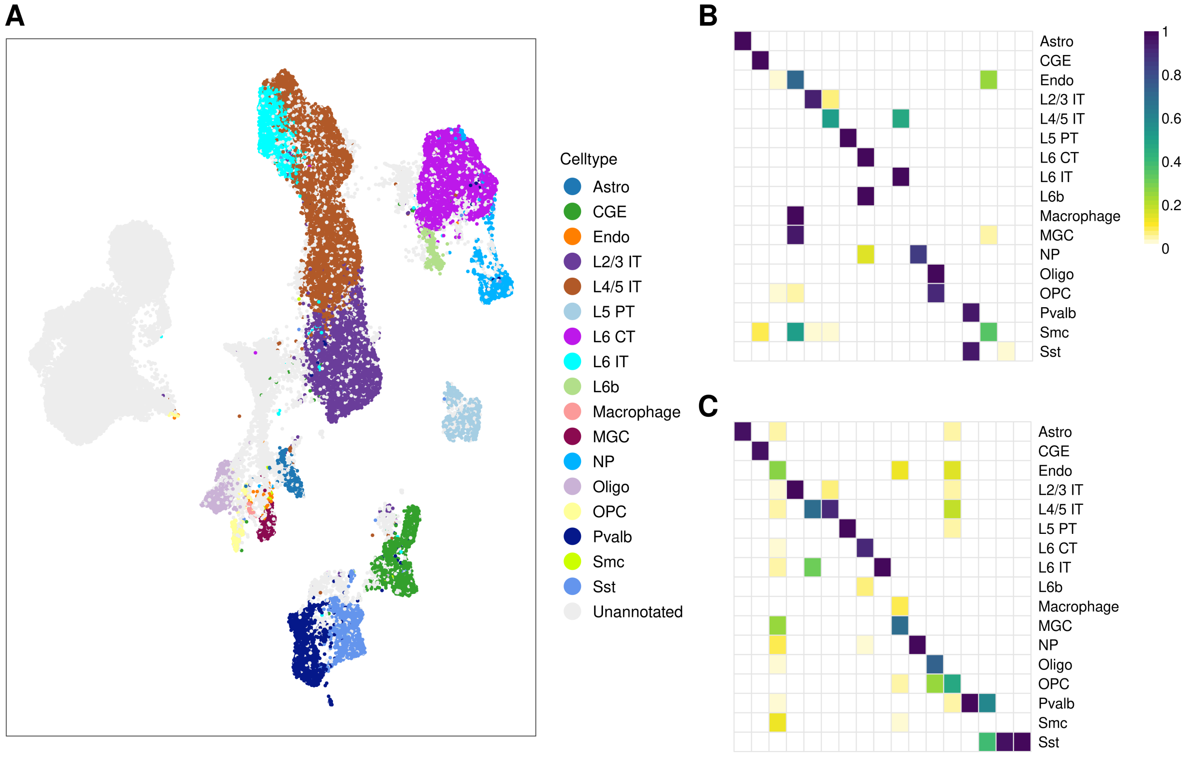


Supplementary Figure 9: `Cobolt` **results on the mouse cortex data integration with no filtering on cells and minimal filtering on features.** Genes and peaks with total counts larger than 10 and non-zero counts larger than 5 are retained in the model. A) UMAP visualization colored by cell type annotation. B, C) Concordance matrices between clustering results and annotations, scaled to sum 1 by (B) rows and (C) columns.

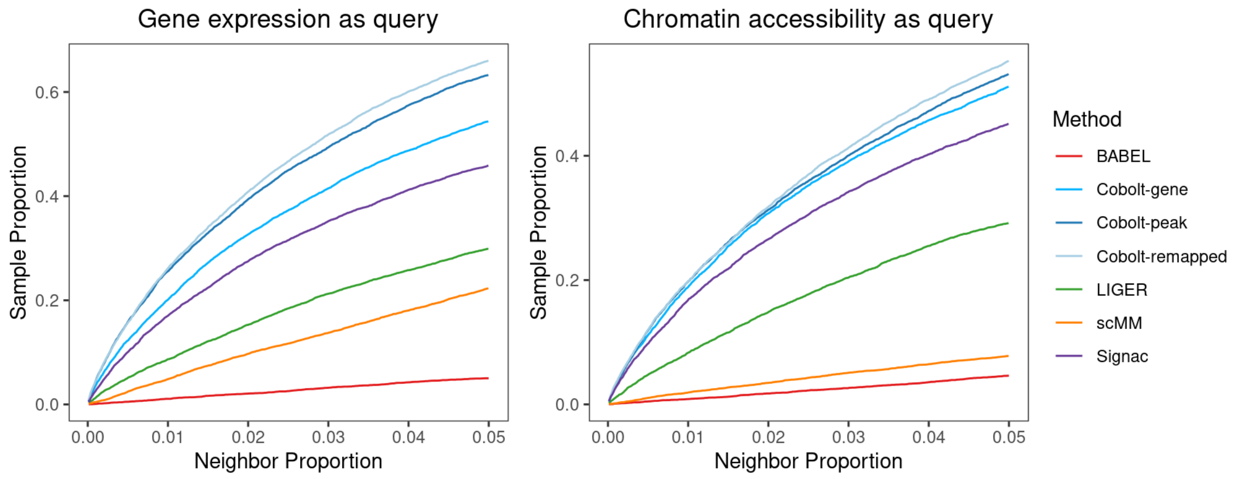


Supplementary Figure 10: **Full results of Figure 5A in the paper (cortical cells)**. Also included is a comparison of `Cobolt` dimensionality reduction on SNARE-seq data 1) using peaks called on the SNARE-seq (`Cobolt`-peak), 2) mapping counts to the peaks called on the MOp scATAC-seq (`Cobolt`-remapped), and 3) summarizing chromatin accessibility counts to gene regions (`Cobolt`-gene). No significant performance loss is observed for `Cobolt`-remapped compared to `Cobolt`-peak, indicating our strategy of mapping data to peaks called on similar datasets a simple but effective alternative in integration analysis. Using peaks vastly improved the performance relative to using the gene activities summarization (`Cobolt`-gene), suggesting that there is a loss of information in using gene summaries, such as distant binding sites not included in the gene summarization.

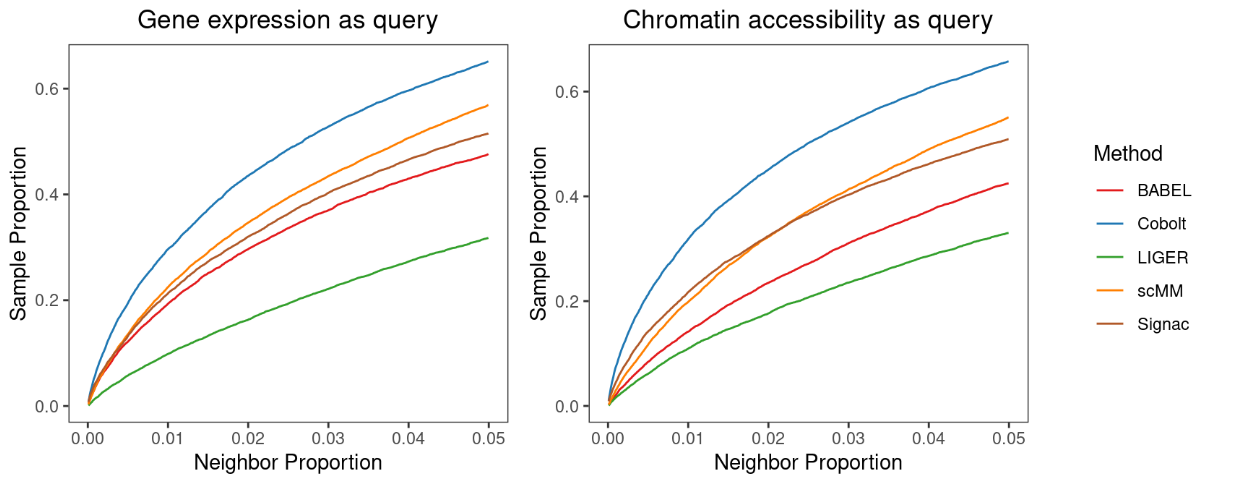


Supplementary Figure 11: Full results of Figure 5B in the paper (PBMC cells).

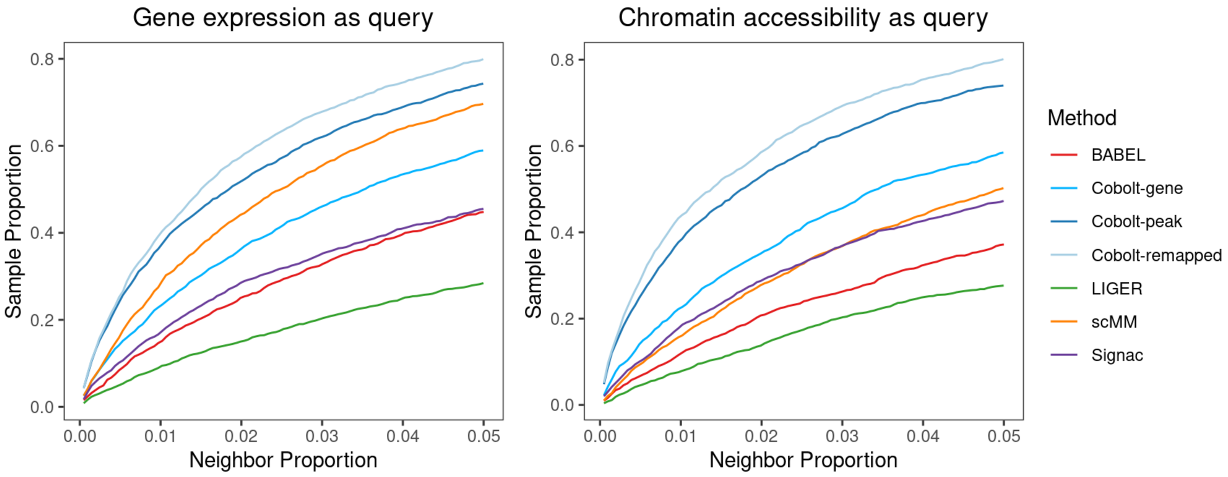


Supplementary Figure 12: An analysis as in Figure 5A in the paper, with a larger percentage of cells used as paired data in the training of scMM, BABEL, and Cobolt (80% rather than 20%). The x-axis shows the number of neighbors considered (k) as a proportion of the total testing sample size. The y-axis shows the proportion of cells whose paired data are within their k-nearest neighbors in the other modality.

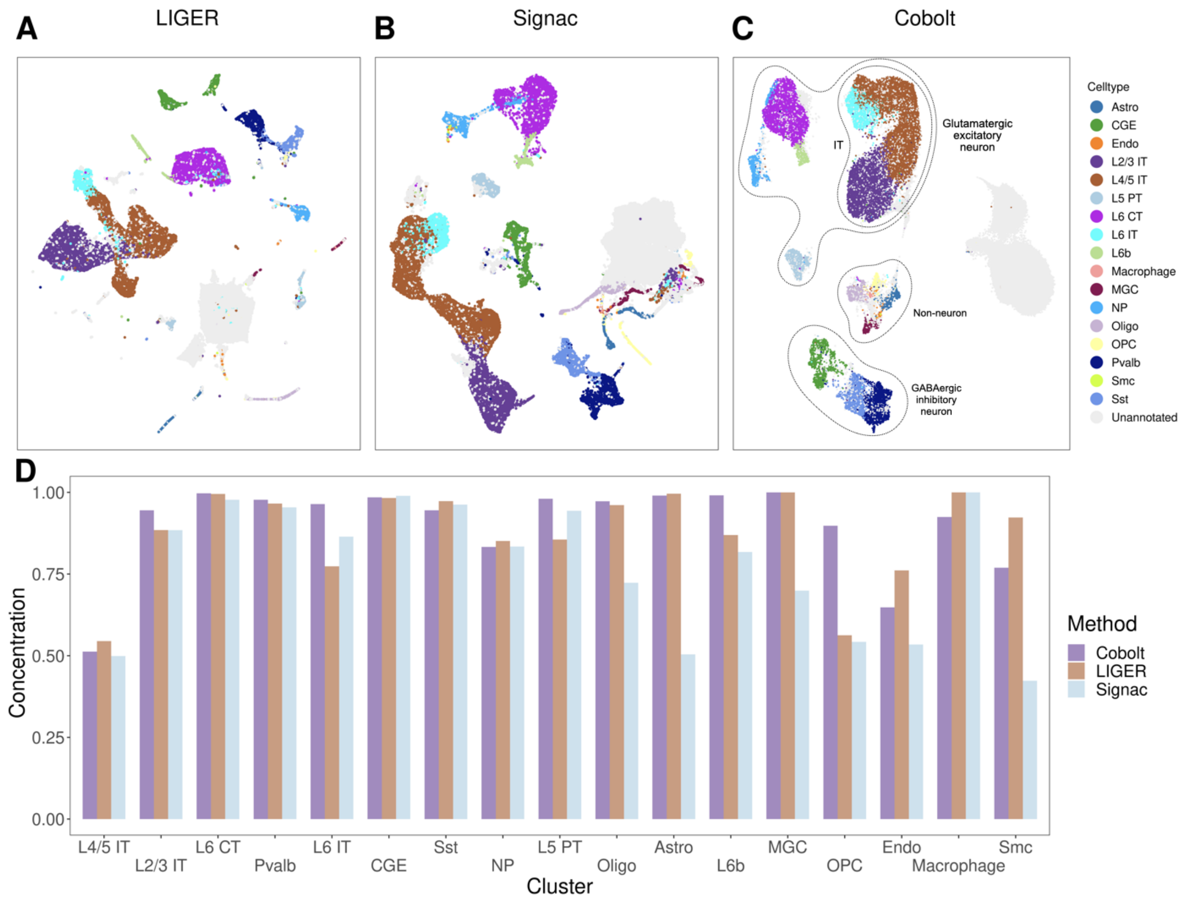


Supplementary Figure 13: A, B, C) UMAP visualizations generated by (A) LIGER, (B) Signac, and (C) `Cobolt` aligning MOp scATAC-seq and scRNA-seq, colored by cell type annotations. The three major classes of cell types—GABAergic inhibitory neurons, glutamatergic excitatory neurons, and non-neurons—are indicated on the `Cobolt` UMAP plot. D) Barplot of maximum overlap proportion between identified clusters and each cell type. For the complete concordance between clustering results and *a priori* cell type labels, see Supplementary Figure 14).

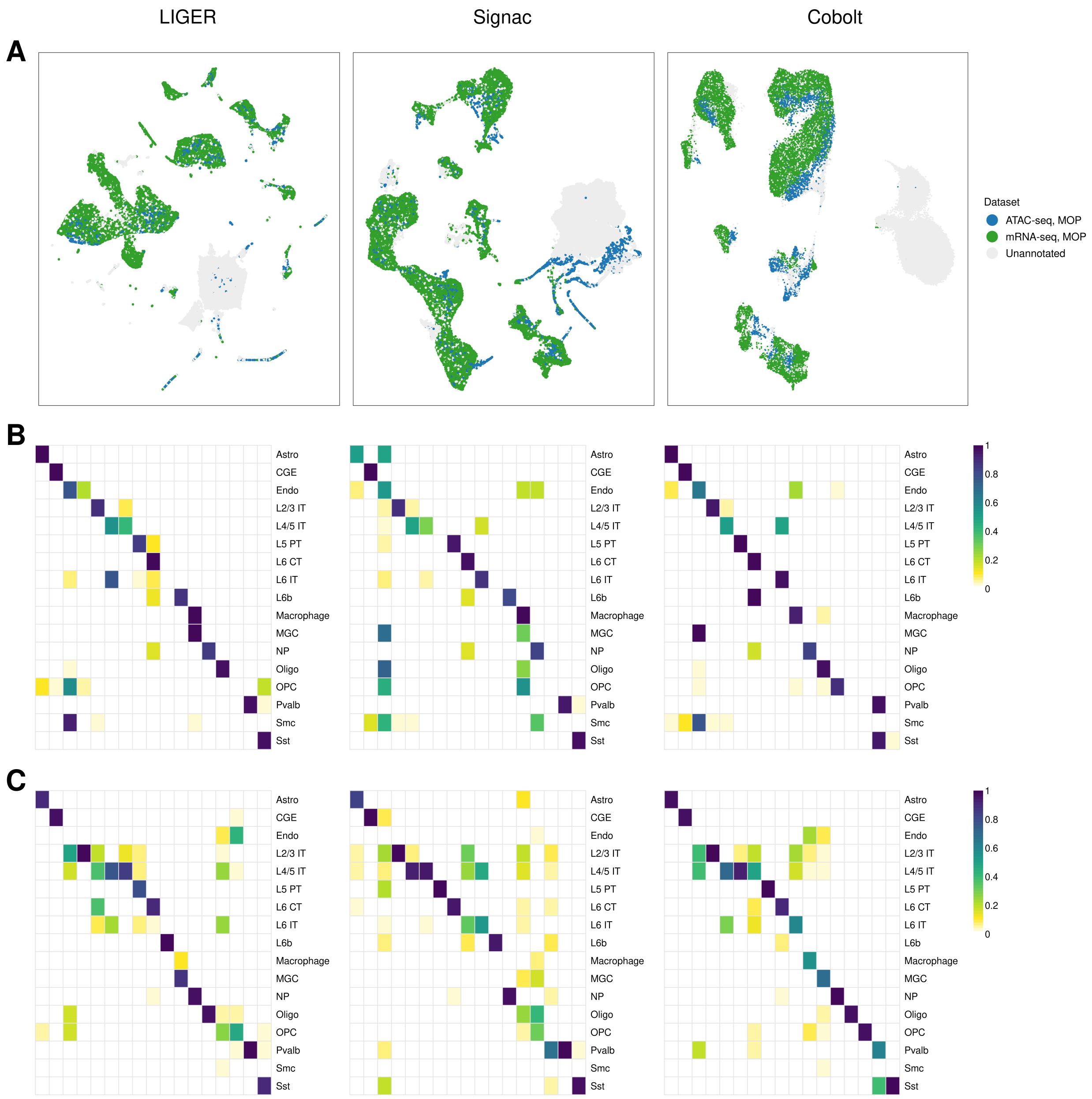


Supplementary Figure 14: A) UMAP visualizations generated by LIGER, Signac, and `Cobolt` for MOp scATAC-seq and scRNA-seq, colored by datasets. B, C) Concordance matrices between clustering results and annotations, scaled to sum 1 by (B) rows and (C) columns.

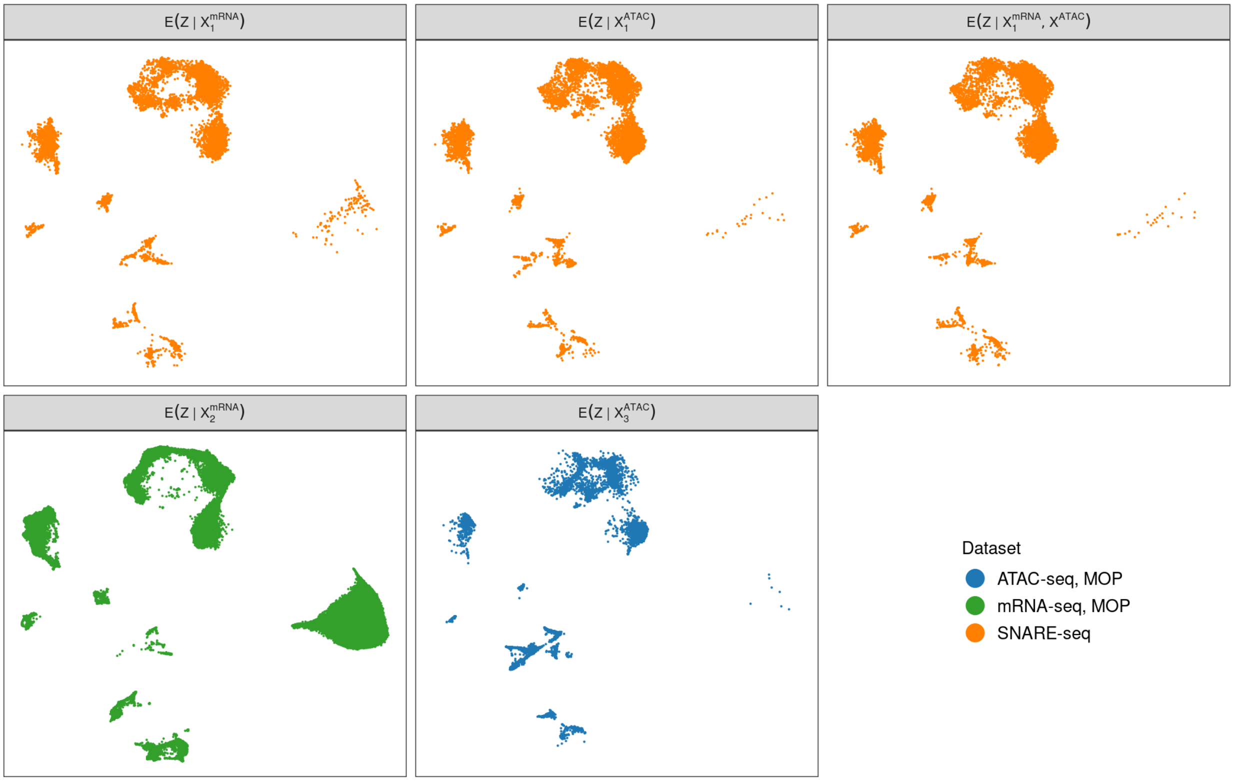


Supplementary Figure 15: **Estimates of the posterior means *without* missing modality correction.** The upper panel shows the UMAP dimensionality reduction plots of the posterior means $\hat{Z}_1 = E(Z|X_1^{mRNA}, X_1^{ATAC})$, $\hat{Z}_1^{mRNA} = E(Z|X_1^{mRNA})$, and $\hat{Z}_1^{ATAC} = E(Z|X_1^{ATAC})$ estimated for joint cells. The lower panel shows $\hat{Z}_2^{mRNA} = E(Z|X_2^{mRNA})$ and $\hat{Z}_3^{ATAC} = E(Z|X_3^{ATAC})$ estimated for single-modality cells. Other than the unannotated cells from MOp scRNA-seq data, cell populations from different datasets are well-aligned. We noticed slight distributional shifts between $\hat{Z}_1$, $\hat{Z}_1^{mRNA}$, and $\hat{Z}_1^{ATAC}$ in the upper panel, motivating additional correction procedures.

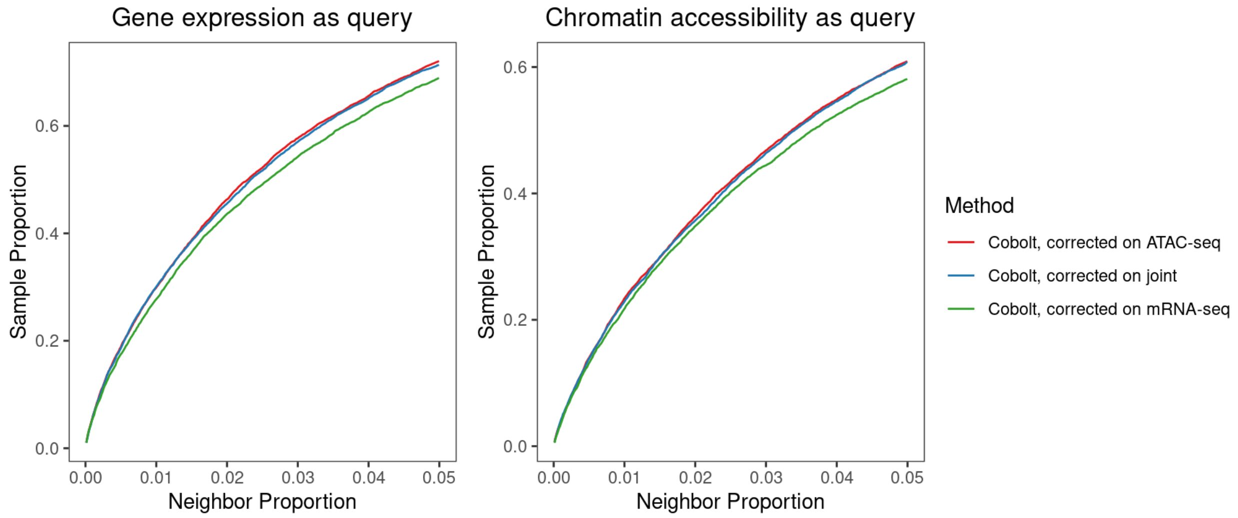


Supplementary Figure 16: **Effect of Prediction Space.** `Cobolt` was applied on the SNARE-seq data sets (similar to that of the Figure 5A) predicting coordinates into the ATAC-seq space $E(Z|X^{ATAC})$ (red), mRNA-seq space $E(Z|X^{mRNA})$ (green), or the joint space $E(Z|X_1^{mRNA}, X_1^{ATAC})$ (blue). Little difference is seen in the quality of the predictions in these different spaces.

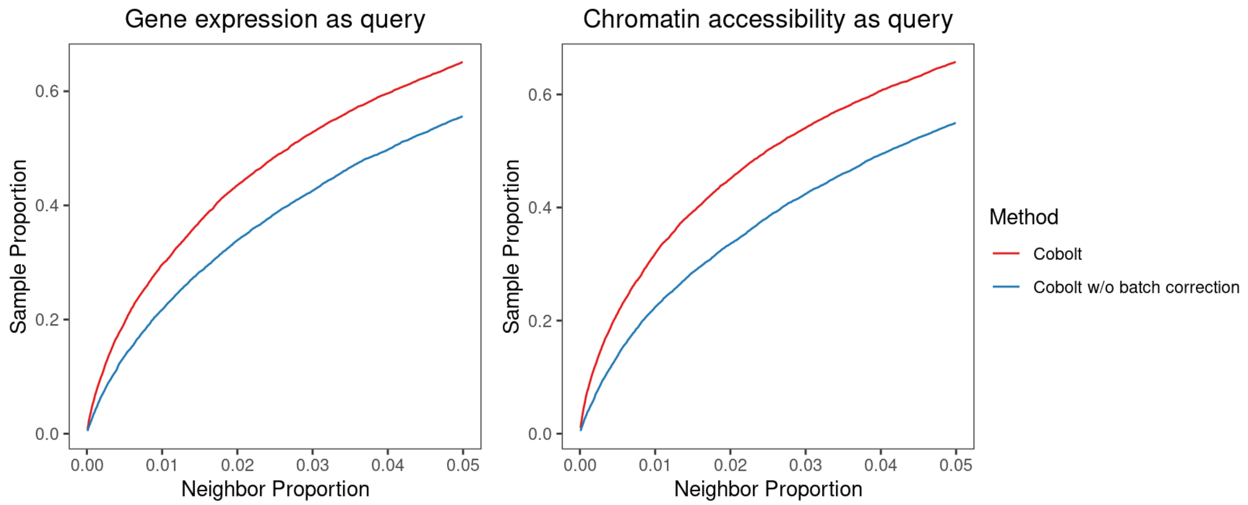


Supplementary Figure 17: **Effect of Batch Correction.** The effect of batch correction of the `Cobolt` model on the results. `Cobolt` was applied on the two 10X multiome data (similar to that of the Figure 5B) with and without the batch correction. The batch correction terms improve the performance as indicated by the closer distance between paired testing cells.

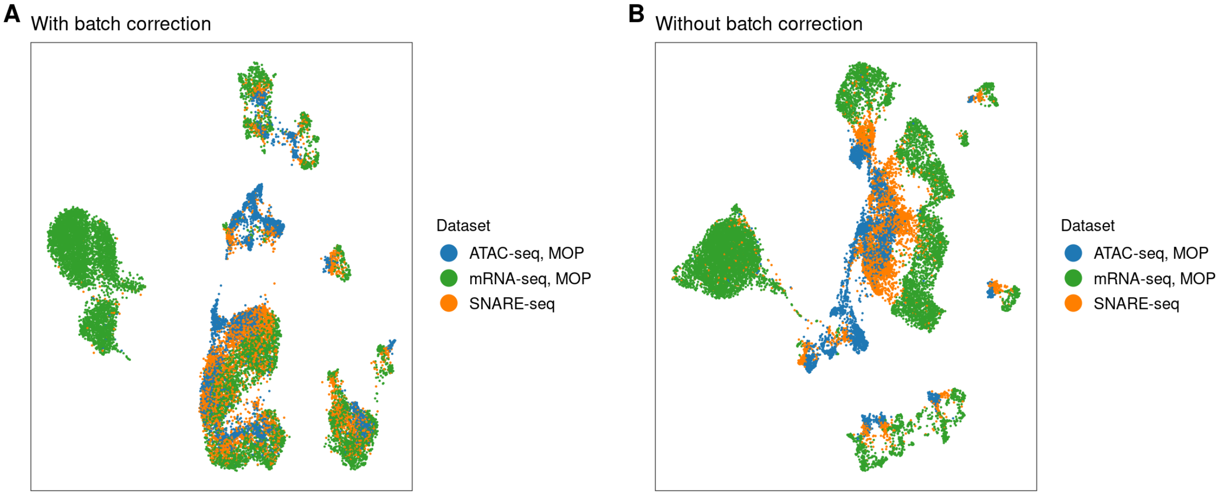


Supplementary Figure 18: **`Cobolt` on SNARE-seq data, with and without batch correction.** UMAP visualizations generated by applying `Cobolt` to the SNARE-seq data (A) with batch correction and (B) without batch correction.
