## Supplementary Text for "Cobolt: Joint analysis of multimodal single-cell sequencing data"

### Supplementary Materials

#### Model Inference

**The Objective Function** To estimate the posterior distribution $p(z|x)$ of the latent variable $z$ and the parameters, we follow the VAE approach to maximize the evidence lower bound (ELBO):

$$\mathrm{ELBO}(x) = \mathbf{E}_{q_\phi(z|x)}[\log p_\psi(x|z)] - \gamma\, \mathrm{KL}(q_\phi(z|x), p(z)),$$

where the encoder $q_\phi(z|x)$ is parameterized by neural networks and serves as an approximation to actual posterior distribution. The decoder $p_\psi(x|z)$ follows our model for single modality, which we will describe in detail later.

The above equation gives the objective function when all the modalities are observed. In case of missing modalities, our conditional independence assumption allows us to obtain a partial ELBO. Consider a subset of modalities $\mathcal{A} \subset \{1, \cdots, M\}$ and data $x^{\mathcal{A}} = \{x^{(i)} | i \in \mathcal{A}\}$, its log-likehood given the latent variable is $\log p_\psi(x^{\mathcal{A}}|z) = \sum_{i\in\mathcal{A}} \log p_\psi(x^{(i)}|z)$. The corresponding ELBO can be written as

$$\mathrm{ELBO}(x^{\mathcal{A}}) = \mathbf{E}_{q_\phi(z|x^{\mathcal{A}})}\left[\sum_{i\in\mathcal{A}} \eta_i \log p_\psi(x^{(i)}|z)\right] - \gamma\, \mathrm{KL}\left(q_\phi(z|x^{\mathcal{A}}), p(z)\right), \tag{1}$$

where $\eta_i$ are hyperparameters allowing different weights for different modalities.

In actual training, even when all modalities are collected, we still subsample the modalities such that posterior likelihoods given different modality subsets are learnt. Denote $\mathcal{S}_c$ as the modalities collected for cell $c$ and $x_c = \{x^{(i)} | i \in \mathcal{S}_c\}$ as the data, our final objective function is given by

$$\sum_{c=1}^{N} \left[\sum_{\mathcal{A}\subseteq\mathcal{S}_c} \lambda_{\mathcal{A}}\, \mathrm{ELBO}(x_c^{\mathcal{A}})\right], \tag{2}$$

where $\lambda_{\mathcal{A}}$'s are hyperparameters allowing different weightings for the ELBO terms with different modalities.

**Prior Distribution** We use Gaussian distribution for the prior $p(z)$. A transformed vector $\theta = \sigma(z)$ is then calculated as the mixing probability of our latent model (Figure 1) for modeling single modalities, where $\sigma$ is the softmax transformation. $\theta$ therefore follows a logistic normal distribution, which is commonly used as an approximation to a Dirichlet distribution. To approximate $Dirichlet(\theta|\xi)$, the parameters of the logistic normal $\mathcal{LN}(\theta|\mu_0(\xi), \Sigma_0(\xi))$ are calculated as a function of $\xi$,

$$\begin{aligned}\mu_0(\xi)_k &= \log \xi_k - \frac{1}{K}\sum_j \log \xi_j,\\ \Sigma_0(\xi)_{kk} &= \frac{1}{\xi_k}\left(1 - \frac{2}{K}\right) + \frac{1}{K^2}\sum_j \frac{1}{\xi_j}.\end{aligned} \tag{3}$$

Therefore, given the hyperparameter $\xi$, we use the Gaussian distribution $\mathcal{N}(z|\mu_0(\xi), \Sigma_0(\xi))$ for the prior $p(z)$.

**Decoders** For cell $c$ in modality $i$, our generative likelihood $p(x_c^{(i)}|\theta_c, B^{(i)}, \alpha_{0,:}^{(i)}, \alpha_{1,:}^{(i)})$ follows multinomial distribution with probability

$$\pi_c^{(i)} = \sigma\big(\operatorname{diag}(\alpha_{1,\ell_c}^{(i)})\sigma(B^{(i)})\theta_c + \alpha_{0,\ell_c}^{(i)}\big), \tag{4}$$

where $\ell_c$ indicates the batch of cell $c$ and $\alpha_{0,\ell_c}^{(i)}, \alpha_{1,\ell_c}^{(i)} \in \mathbf{R}^{d_i}$, and $B^{(i)} \in \mathbf{R}^{d_i \times K}$ are parameters to be estimated that are shared across samples. $\sigma(B^{(i)})$ gives the probability vectors for each latent category. $\alpha_{0,\ell_c}^{(i)}$ and $\alpha_{1,\ell_c}^{(i)}$ therefore provides scaling and shifting for adjusting dataset- or platform-specific differences in feature counts. Notice that Equation 4 works without the softmax transformation on $B^{(i)}$, which gives us another version of the model

$$\pi_c^{(i)} = \sigma\big(\operatorname{diag}(\alpha_{0,\ell_c}^{(i)})B^{(i)}\theta_c + \alpha_{0,\ell_c}^{(i)}\big). \tag{5}$$

In practice, we find 5 works slightly better than 4. The results we present in this paper are based on 5.

In summary, we write the joint likelihood of the latent variable $z_c$ and the data $x_c$ of cell $c$ as,

$$p(x_c^{\mathcal{S}}, z_c|B^{\mathcal{S}}, \alpha_{0,:}^{\mathcal{S}}, \alpha_{1,:}^{\mathcal{S}}) = p(z_c)\prod_{i\in\mathcal{S}} p(x_c^{(i)}|\theta_c = \sigma(z_c), B^{(i)}, \alpha_{0,:}^{(i)}, \alpha_{1,:}^{(i)}), \tag{6}$$

where $p(x_c^{(i)}|\theta_c, B^{(i)}, \alpha_{0,:}^{(i)}, \alpha_{1,:}^{(i)})$ follows multinomial distribution with probability vector given by Equation 5. Supplementary Figure 1 (Additional File 1) shows a graphical model representation of this data generation process.

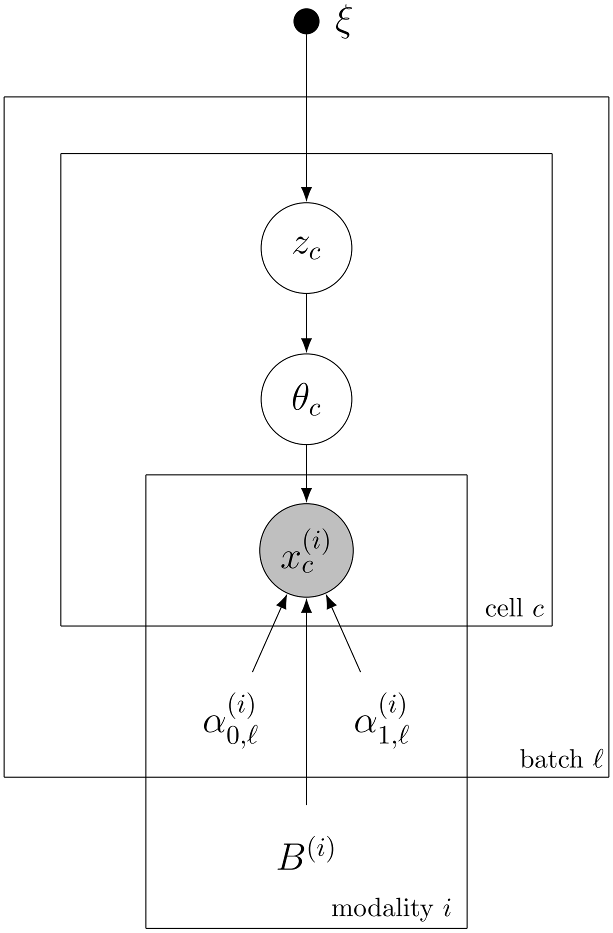


Figure 1: Graphical model representation of the generative distribution of `Cobolt`.

**Encoders** We would like to obtain an approximation $q(z|x^{\mathcal{A}})$ of the posterior distribution $p(z|x^{\mathcal{A}})$ given any collection of modalities $x^{\mathcal{A}}$. To do this, we first write the true posterior as

$$p(z|x^{\mathcal{A}}) \propto p(z) \prod_{i \in \mathcal{A}} \frac{p(z|x^{(i)})}{p(z)}.$$

We then approximate $\frac{p(z|x^{(i)})}{p(z)}$ with Gaussian $g(z|x^{(i)}) := \mathcal{N}(\tilde{\mu}^{(i)}, \tilde{\Sigma}^{(i)})$, where $\tilde{\mu}^{(i)}$ and $\tilde{\Sigma}^{(i)}$ are estimated by neural networks. Here we use the property that the product of two multivariate Gaussian densities is proportional to a Gaussian density. $q_\phi(z|x^{\mathcal{A}}) \propto p(z) \prod_{i \in \mathcal{A}} g(z|x^{(i)})$ is still Gaussian whose parameters can be analytically written out,

$$\begin{aligned} \Sigma^{\mathcal{A}} &= \Big(\tilde{\Sigma}_0^{-1} + \sum_{i \in \mathcal{A}} (\tilde{\Sigma}^{(i)})^{-1}\Big)^{-1}, \\ \mu^{\mathcal{A}} &= \Sigma^{\mathcal{A}} \Big(\tilde{\Sigma}_0^{-1} \tilde{\mu}_0 + \sum_{i \in \mathcal{A}} (\tilde{\Sigma}^{(i)})^{-1} \tilde{\mu}^{(i)}\Big). \end{aligned} \tag{7}$$

With this trick, instead of using $2^M - 1$ neural networks to estimate all $\Sigma^{\mathcal{A}}$ and $\mu^{\mathcal{A}}$ individually, our model uses one network for $\tilde{\mu}^{(i)}$ and $\tilde{\Sigma}^{(i)}$ separately for each modality $i$, resulting in a total of $2M$ networks. The variational posteriors $q_\phi(z|x^{\mathcal{A}})$ are calculated using $\tilde{\mu}^{(i)}$ and $\tilde{\Sigma}^{(i)}$. The variational posterior $q_\phi(\theta|x^{\mathcal{A}})$ is then defined to be logistic normal $\mathcal{LN}(\mu^{\mathcal{A}}, \Sigma^{\mathcal{A}})$.
